# Center-surround interactions underlie bipolar cell motion sensing in the mouse retina

**DOI:** 10.1101/2021.05.31.446404

**Authors:** Sarah Strauss, Maria M Korympidou, Yanli Ran, Katrin Franke, Timm Schubert, Tom Baden, Philipp Berens, Thomas Euler, Anna L Vlasits

## Abstract

Motion is a critical aspect of vision. We studied the representation of motion in mouse retinal bipolar cells and found, surprisingly, that some bipolar cells possess motion-sensing capabilities that rely on their center-surround receptive fields. Using a glutamate sensor, we directly observed motion-sensitive bipolar cell synaptic output, which was strongest for local motion and dependent on the motion’s origin. We characterized bipolar cell receptive fields and found that there are motion and non-motion sensitive bipolar cell types, the majority being motion sensitive. Next, we used these bipolar cell receptive fields along with connectomics to design biophysical models of downstream cells. The models and experiments demonstrated that bipolar cells pass motion-sensitive excitation to starburst amacrine cells through direction-specific signals mediated by bipolar cells’ center-surround receptive field structure. As bipolar cells provide excitation to most amacrine and ganglion cells, their motion sensitivity may contribute to motion processing throughout the visual system.

## Introduction

Local motion sensing is of paramount importance for sighted animals, enabling them to detect and capture prey (1–4), as well as to avoid predators (5–9). In mammals, multiple features related to motion sensing are first extracted from the visual scene by the retina (10,11). These features include the direction of motion (12, 13), looming motion (14), and differential motion (15), and can be used, for instance, to filter the local motion of objects from the global motion caused by body, head, and eye movements. The stages at which motion is extracted in the retinal circuitry and the mechanisms of motion-related feature detection are key to understanding these processes.

Motion features are most likely to be computed in the inner retina. There, 14 types of bipolar cells (BCs, (16–20), or 15 if the so-called GluMI is included (21)), receive input from photoreceptors. BCs provide excitatory glutamatergic input to a large diversity of amacrine cells (ACs), which are a class of inhibitory interneurons (reviewed in (22)), and retinal ganglion cells (RGCs), which are the output neurons of the retina (reviewed in (23, 24)). Although BCs represent the first stage in the retina where visual signals diverge into parallel channels, motion detection has not yet been found to be implemented at the BC level.

For instance, one well-studied motion detection circuit in the retina is the direction selectivity (DS) circuit, where the BCs’ role in the motion computation remains intensely debated. One key element in this DS circuit is the starburst amacrine cell (SAC), which exhibits DS for motion at the level of individual neurites (25), providing asymmetric inhibition to direction-selective RGCs during motion in one direction (26–30). The role of BCs in this DS circuit has been a matter of intense scrutiny, with a variety of studies having provided evidence supporting an important role for BCs in the motion computation by SACs ((19, 31, 32), but also see (33, 34)), or by direction selective RGCs (35, 36). More specifically, it was suggested that distinct BC types with different glutamate release kinetics (16, 37, 38) provide spatially offset inputs on postsynaptic SAC dendrites (“space-time” wiring, (31)), enhancing the preferred direction response. While voltage-clamp recordings in the rabbit retina implicate some directional tuning in BCs (39–41), BC Ca^2+^ signals and glutamate release have been observed to respond symmetrically to motion stimuli (42–45) (but see (46)). Therefore, BCs have not been considered direction-tuned cells themselves and their exact role in the DS circuit remains debated.

At the same time, BCs exhibit a basic receptive field (RF) feature that could support motion detection: their center-surround antagonism (47–50). Center-surround antagonism refers to the fact that BCs prefer opposite polarity stimuli in the center of their RFs vs. the surround. So-called On BCs depolarize and release more glutamate to light increments in the center (a light turning “On”) and light decrements in the surround, while Off BCs have the opposite preference.

Antagonistic interactions between RF center and surround enrich the BC types’ functional diversity. BC types possess differences in the size and strength of center and surround as well as in the temporal relationship between center and surround responses (16, 51). These interactions are partially established by horizontal cells in the outer plexiform layer (reviewed in (52)), but importantly shaped further by ACs (16, 53). More than 50 years ago, it was hypothesized that the interplay between spatially and temporally offset excitation and inhibition establishes retinal motion detectors (54). Yet, the role of these antagonistic center-surround RF interactions in local motion detection has not been extensively explored (55, 56).

Here, we studied the local motion sensing properties of BCs throughout the inner retina by measuring BC output using a fluorescent glutamate sensor during visual stimulation. Surprisingly, we found that some BCs exhibit a sensitivity to motion direction conditioned on the origin of motion. To explore this further, we characterized the center-surround RFs of BCs across the inner plexiform layer (IPL) and uncovered diversity in their RF properties for motion sensing, which confers looming and direction sensitivity to a subpopulation of BC types. We explored the implications of these motion sensing properties for downstream retinal processing in SACs by constructing biophysical models of SAC dendrites with anatomically and spatio-temporally precise input from BCs. We found that the SAC inherits directionally tuned input from BCs during local motion stimulation and this DS is diminished by *in-silico* removal of the BCs’ RF surrounds. Last, we verified our *in-silico* findings experimentally by measuring glutamate release onto SACs and Ca^2+^ dynamics in SAC dendrites. Our findings suggest that BCs produce direction selective signals for motion originating in their RF centers, and that these signals can play a role in the computation of local motion direction in SACs. Given the central role of BCs in retinal signaling, our findings suggest that BCs may play a key role in many motion computations throughout the retina.

## Results

### Complex bipolar cell glutamate release in response to local motion

To observe how BCs respond to small, locally moving stimuli, we performed 2-photon imaging of a glutamate sensor, iGluSnFR (16, 45) expressed throughout the neurons of the IPL (**Fig. 1A**). We began by imaging at relatively low spatial resolution to observe glutamate release dynamics over a large field of view (FOV, ~200 μm) during visual motion stimulation. First, we presented a small bright moving bar (20 × 40 μm) that traversed a distance of 100 μm, corresponding to roughly 3.3°of visual angle (57) spanning the width of 2-4 BC RFs, in two opposite directions while imaging in the On layer of the IPL (**Fig. 1B**). Glutamate signals from this stimulus were complex, with changes in glutamate release occurring throughout the FOV, well beyond the bounds of the stimulus and its trajectory (**Fig. 1B**). Surprisingly, we found that the response amplitude was motion direction sensitive in some regions of the FOV. Specifically, the region of interest (ROI) in which the stimulus originated (Box 1, **Fig. 1C**) exhibited more glutamate release for motion out of the FOV compared to motion into the FOV. In a nearby ROI (Box 2) the glutamate release appeared symmetric between motion directions.

**Fig. 1.**
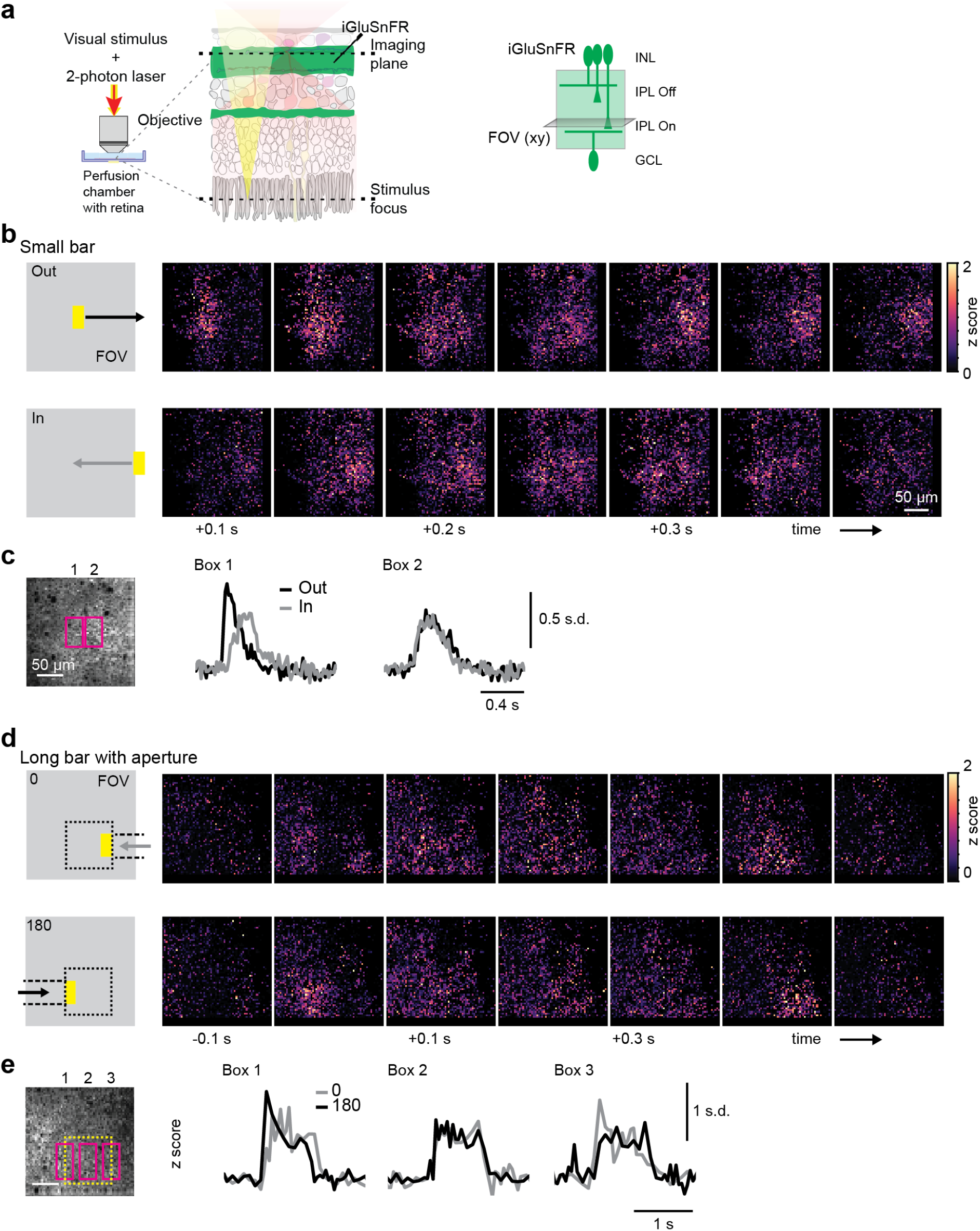
Complex bipolar cell glutamate release in response to moving stimuli. **(a)** Left: Experimental setup showing objective and retina, with visual stimulus (yellow) and 2-photon laser (red). Right: iGluSnFR is ubiquitously expressed in retinal neurons, including in the cells of the IPL (green region). INL, inner nuclear layer; IPL, inner plexiform layer; GCL, ganglion cell layer; FOV, field of view. **(b)** Left, “Small bar” stimulus (20 x 40 μm light rectangle on dark background) moving at 500 μm/s over a distance of 100 μm, beginning either in the center of the FOV or just outside the FOV. Right, montage of the average z-scored fluorescence response of glutamate sensor iGluSnFR during stimulation in each direction. **(c)** Left, the average iGluSnFR fluorescence during stimulation, showing two ROIs used to measure fluorescence responses. Right, mean binned fluorescence in response to each stimulus condition for the pixels in each ROI. **(d)** Left, “long bar with aperture” (40 × 385 μm, appearing only in the dotted square) moving at 500 μm/s in two directions (0° vs. 180°). Right, montage as in (b). **(e)** Average responses in three regions, as in (c).

We next sought to determine whether this preference for motion origin was stimulus type-specific. We therefore displayed a widely-used type of moving bar stimulus, in which a thin, long bar moves across a patch of retina in two directions (i.e. (39, 58)). To capture the appearance of this motion in our FOV, we restricted the area through which the bar moved to a rectangle smaller than our imaging FOV (dotted box, **Fig. 1D**). Here again, we found that regions where the motion originated (Box 1 and 3, **Fig. 1E**) exhibited a preference for the motion direction that originated within that area, while a region in the center of the bar’s motion trajectory exhibited symmetric responses (Box 2). Thus, BCs appear to signal the origin of moving stimuli.

### Bipolar cell glutamate release is sensitive to motion origin

To further explore BC motion sensitivity, we sought to measure whether individual BC terminals exhibit a preference for motion origin. We performed iGluSnFR imaging at higher spatial resolution in the On layer of the IPL (**Fig. 2A**) and presented moving stimuli originating inside the FOV and moving out (“out”) or outside of the FOV and moving in (“in”) and traversing different distances (**Fig. 2B**). To better capture the activity of small, noisy ROIs that were the size of BC terminals (16) (see Methods for details), we used Gaussian Process modeling to infer the mean and standard deviation (s.d.) of individual ROI responses to each stimulus condition (59) (Figure S2a). We observed that many ROIs exhibit strong glutamate release to motion originating in the FOV (“100 out” stimulus), and that ROIs preferred this stimulus to motion in the opposite direction (“100 in”) (**Fig. 2C**). We calculated the extent of this preference (d-prime, *d*′) to examine stimulus preference across the FOV (**Fig. 2C-D**). We found that the preference for motion origin was restricted to a small region of about the size of a BC’s RF center, and that there was no direction preference when the motion originated outside of the FOV (**Fig. 2E**, data from n=4,056 ROIs/ 7 fields/ 3 mice). In addition, we found that within the area where the motion originated, the *d*′ across ROIs was significantly shifted toward positive values, signifying a preference for motion going out of the FOV (**Fig. 2F-G**, 100 μm, *d*′ = 41.0 ± 54.1; 150 μm, *d*′ = −2.4 ±26.7; 300 μm, *d*′ = 4.4 ±30.0; *p* < 0.01, Wilcoxon signed-rank test, n=641 ROIs/ 7 fields/ 3 mice). These results suggest that at least some BC types are highly sensitive to the precise origin of a motion stimulus.

**Fig. 2.**
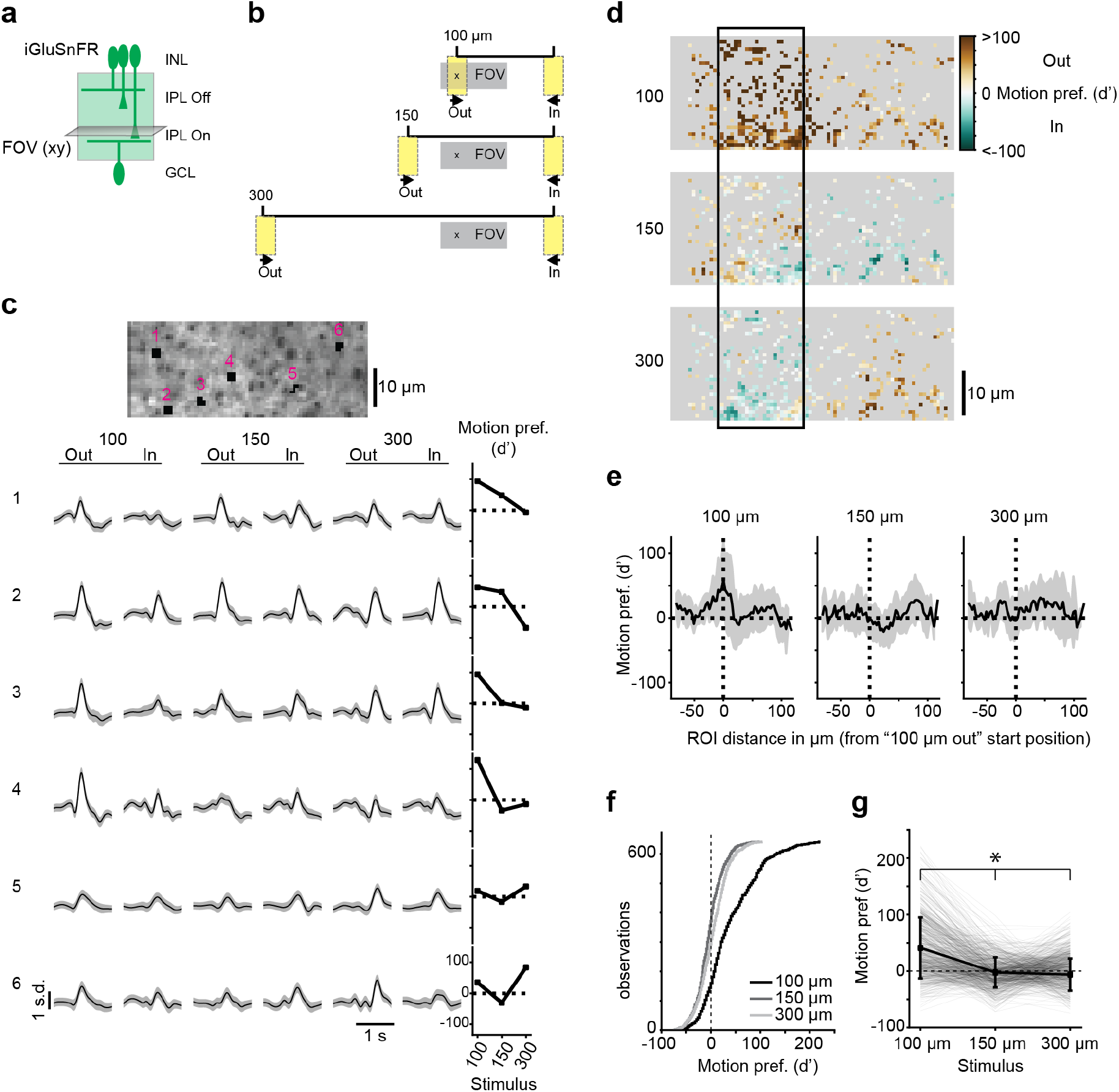
Bipolar cell glutamate release is sensitive to motion origin. **(a)** iGluSnFR is ubiquitously expressed in retinal neurons, including in the cells of the IPL (green region). INL, inner nuclear layer; IPL, inner plexiform layer; GCL, ganglion cell layer; FOV, field of view. **(b)** Moving bars (20 x 40 μm) presented to the retina traveling in two directions (out vs. in) and traversing 3 distances (100, 150, 300 μm). All objects to scale. **(c)** Example ROIs (black regions, numbered) overlaid with s.d. of the imaged field and their responses to the stimuli in (b) as predicted using Gaussian Process modeling. Grey shading is 3 s.d. Rightmost column: motion preference (*d*′) for each stimulus travel distance. Positive values represent a preference for motion in the “out” direction. **(d)** The motion preference (*d*′) for all ROIs in example field. The boxed region is the starting position of the “100 out” condition and is analyzed in (f) and (g). **(e)** Motion preference (*d*′) for each stimulus condition as a function of location relative to the “100 μm out” start position (“x” in b). Sample size is n=4,056 ROIs/ 7 fields/ 3 mice. **(f)** Cumulative histogram of motion preference (*d*′) for ROIs located within 10 μm on either side of the “100 μm out” stimulus start position (“x” in b; black rectangle in d). **(g)** Motion preference (*d*′) for each ROI in the population for each stimulus condition. All conditions are significantly different (*p* < 0.01, Wilcoxon signed-rank test). Sample size in (f-g) is 641 ROIs/ 7 fields/3 mice. See also Figure S2a, Figure S2

### Bipolar cells exhibit differing sensitivity to motion

Previous measures of BC response properties suggest that the 14 BC types differ in their spatial and temporal response properties and kinetics (16, 38, 60, 61). These differences could be important for motion sensing. Thus, we sought to determine whether all BC types exhibit sensitivity to motion origin. We used 2-photon volumetric imaging enabled by an electrically-tunable lens (62). This allowed “axial” (x-z) scans and, hence, to image glutamate release across all IPL layers at once. Initial observations of responses to moving bar stimuli suggested that not all BCs are sensitive to motion origin (Figure S2b). To determine the extent of motion sensitivity across BC types, we measured the RF properties of BC glutamate release using a “1D noise” stimulus (**Fig. 3A**) and inferred smooth RFs using a spline-based method (63). In this way, we could observe center-surround RFs from ROIs near the size of individual BC boutons (ROI sizes ~2 μm^2^, see **Methods**), including clear On and Off RFs from their respective IPL strata (**Fig. 3B**) for 3,233 ROIs. We clustered these RFs into groups using a Mixture of Gaussian clustering on features from the RFs as well as each ROI’s IPL depth, and uncovered 13 clusters of BC RFs (**Fig. 3C-E**). Individual clusters contained ROIs stratifying tightly in the IPL (**Fig. 3F**) and most clusters exhibited stereotyped temporal properties of their centers and surrounds within cluster (Figure S3b). We computed the average RF for each cluster and observed that these average RFs had distinct properties, most notably the temporal and spatial characteristics of the surround (**Fig. 3G-H**, see also **Fig. 4**).

**Fig. 3.**
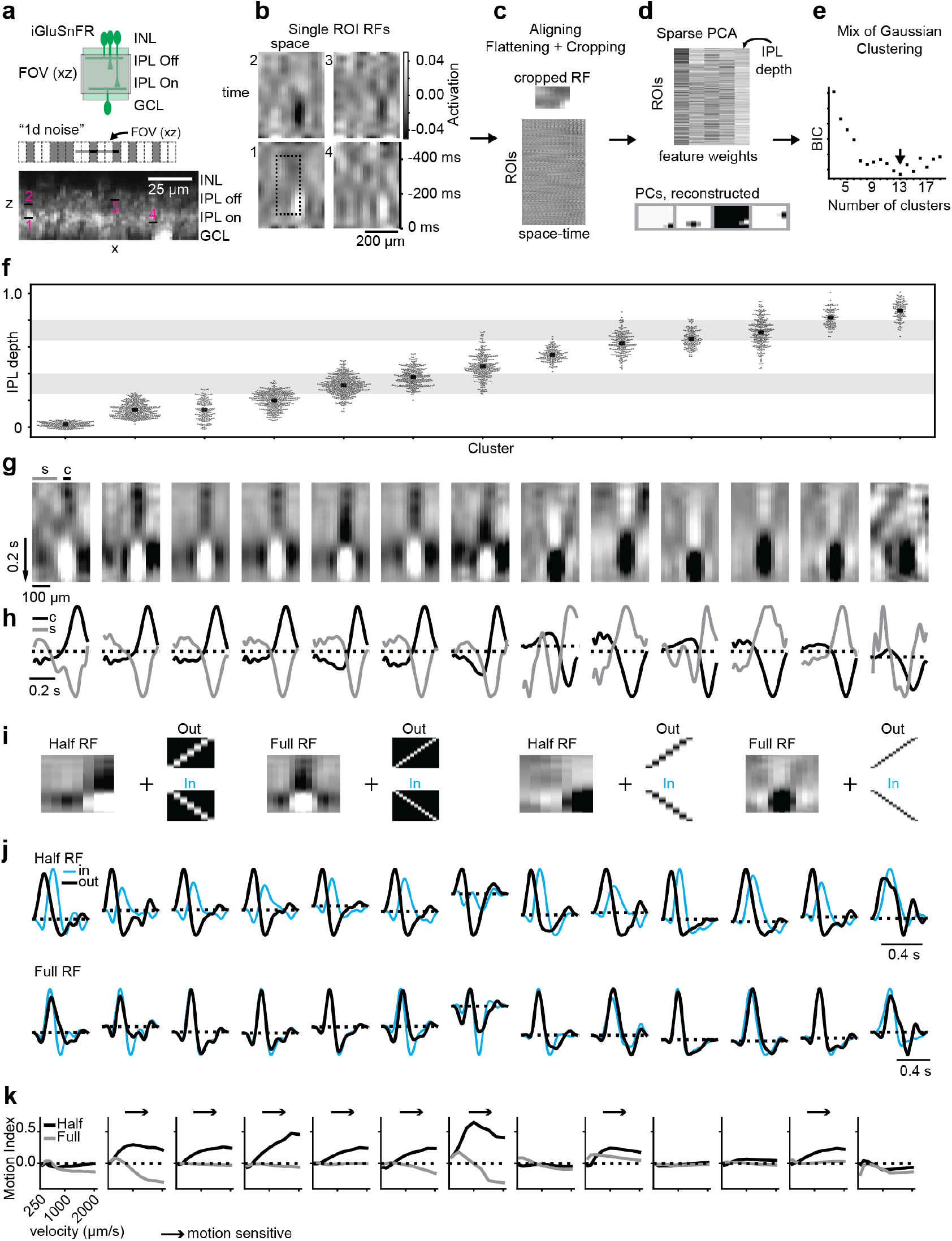
Bipolar cell receptive fields exhibit differing sensitivity to motion. **(a)** Top: iGluSnFR ubiquitously expressed as in Fig. 1. Imaging is performed using an electrically-tunable lens to achieve x-z scanning. INL, inner nuclear layer; IPL, inner plexiform layer; GCL, ganglion cell layer; FOV, field of view. Middle: “1D noise” stimulus consisting of twenty 20×50 μm rectangles switching randomly between black and white at 20 Hz. The relative scale of the x-z scan field (FOV) is shown. Bottom: average of a scan. Black regions/numbers: ROIs analyzed in (b) **(b)** Example RFs from four ROIs from the field in (a, numbers). The top examples are from the Off layer, the bottom examples are from the On layer. Dotted box: the cropped RF used for clustering. **(c)** Top: the cropped RF from dotted box in (b), with the same aspect ratio as the PCs in (d). Bottom: RFs were aligned to the RF center and then all RFs were flattened to 1 dimension and cropped to exclude missing space-time. The final dataset includes RFs from 3,233 ROIs/ 4 fields/ 4 eyes/ 3 mice. **(d)** Top: feature weights for the 4 components from PCA. The fifth feature was the IPL depth of the ROI. Bottom, reconstructed components of the sparse PCA. **(e)** Mixture of Gaussian clustering was performed and the Bayesian information criterion (BIC) was used to select the number of clusters. **(f)** Cluster assignment of each ROI plotted against IPL depth. Clusters were reordered by average IPL depth. Grey regions: approximate ChAT bands (the dendritic plexi of the SACs as an IPL landmark). **(g)** Average RF of each cluster. “c” and “s” show the regions used to calculate the spatial average of the center and surround in (h). **(h)** Average temporal RFs taken from the center (“c”) and surround (“s”) regions indicated in (g), normalized to their respective peaks. **(i)** Example RF showing the cropped RFs (“half” vs. “full”) convolved with the motion stimuli (“out”, black, vs. “in”, cyan) to measure the motion sensing properties of each cluster. **(j)** Modeled responses to motion (velocity 1,000 μm/s) in two directions for each cluster for stimuli passing over the full RF or half of the RF. **(k)** Motion index as a function of velocity for each cluster for the full (grey) vs. half (black) conditions. See also Figure S3a, Figure S3b.

**Fig. 4.**
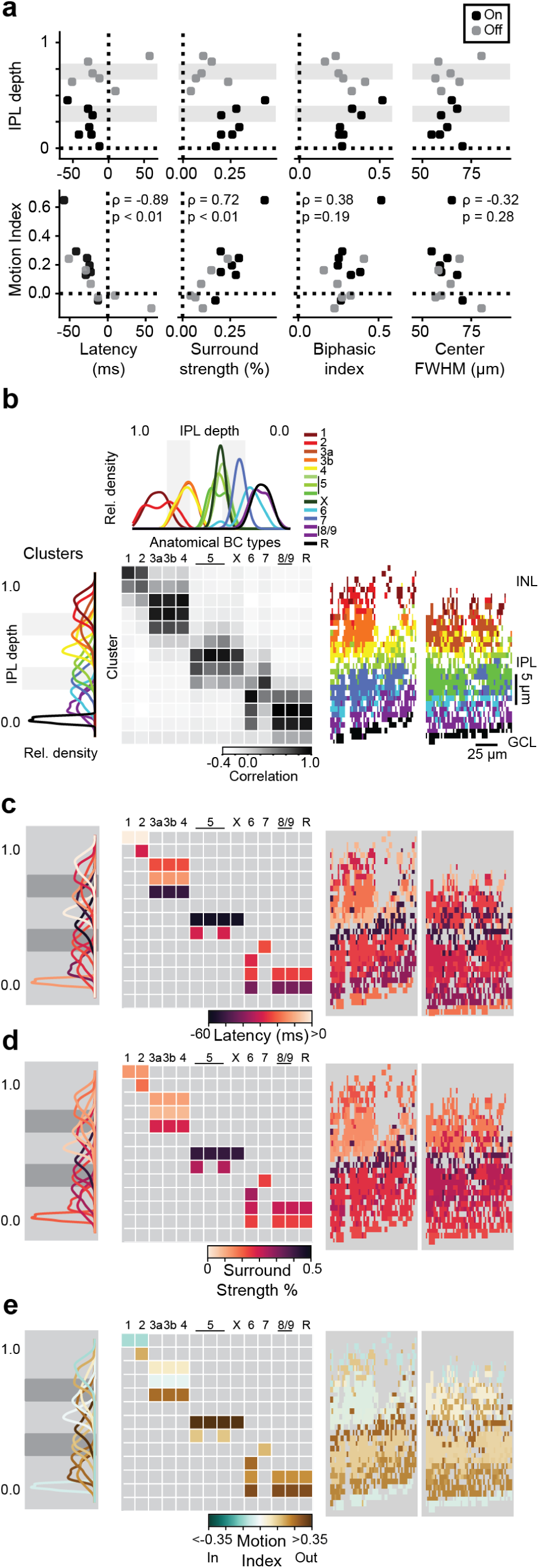
Layer-specific motion sensitivity depends on bipolar cell surround. **(a)** Center-surround properties of BC clusters from Fig. 3 plotted against IPL depth (top) or Motion Index of modeled responses (bottom). Black and gray points represent On and Off-type BCs, respectively; gray shading marks approx. ChAT bands. **(b)** Top: density plot of BC anatomical stratification as reported earlier (18, 19, 31). Left: density plot of BC cluster stratification of ROIs from each cluster in Fig. 3. Colors chosen by likely matches with anatomical stratification. Grey shading marks approx. ChAT bands. Middle: correlation between BC clusters and anatomical types based on their stratification in the IPL. Right: cluster assignment and likely BC type mapped onto pixels from two example imaging fields. **(c)** Left: density plot from (b) color coded by the latency between peak of surround and center responses. Middle: cluster latency assigned to squares with greater than 0.7 correlation based on IPL stratification in (b). Right: cluster latency of surround vs. center mapped onto ROIs from two imaging fields (same fields as (b)). **(d)** Same as (c), but with all panels color coded by the cluster surround strength relative to center strength. **(e)** Same as (c), but with all panels color coded by the cluster Motion Index measured from RF convolution with 1,000 μm/s velocity motion stimulus (**Fig. 3J)**

To evaluate the motion sensing properties of individual BC clusters, we modeled their responses to a moving bar stimulus by convolving the average RF for each cluster with a space-time stimulus image (**Fig. 3I**). To test for a preference for motion origin, we played the stimulus to just one half of the RF, which corresponds to a bar originating in the RF center and moving to the surround, or vice versa for the opposite motion direction. We compared this scenario to the case of a bar moving through the full RF, from surround to center and then surround again. We found that some BC clusters exhibited a preference for motion originating in their RF centers, while others showed no preference for motion originating in the RF center or surround (**Fig. 3J-K**). We modeled these responses across a range of stimulus velocities and measured a motion sensitivity index (Motion Index) for these stimuli across velocities. We found that some BC clusters in both On and Off layers exhibited motion sensitivity across a range of velocities, while other clusters were not motion-sensitive (**Fig. 3K**). In addition, there was very little directional preference for any cluster in response to stimulation across the full RF. We also found that motion origin sensitivity was not limited to the moving bar stimulus, but that motion-sensitive BCs also preferred looming stimuli to receding stimuli (Figure S3a), suggesting that specific BC types might be important for several types of motion sensing tasks that are known to be performed within the retina (4, 6, 10).

### Layer-specific motion sensitivity depends on bipolar cell surround

To determine which features of the BC RFs are important for establishing motion sensitivity, we measured several properties of the cluster RFs and found that longer center-surround latency and stronger surround strength were correlated with increased motion sensitivity (**Fig. 4A**, center-surround latency vs. Motion Index, Spearman correlation *ρ* = −0.89, *p* < 0.01; surround strength vs. Motion Index, *ρ* = 0.72, *p* < 0.01), while properties of the center were not (biphasic index vs. Motion Index, *ρ* = 0.38, *p* = 0.19; center full-width half-max (FWHM) vs. Motion Index, *ρ* = −0.32, *p* = 0.28). These results suggest that BC motion detection operates like a Barlow-Levick detector (54), in which spatial and temporal offset of the inhibitory surround and excitatory center establish sensitivity to motion (see **Fig. 5B**). We confirmed the critical role of the BC RFs’ surround in the modeled motion preferences by decreasing the strength of the surround artificially (Figure S5a). Then, by decomposing the modeled responses into contributions from the center and surround, we observed that the inhibition from the surround is more temporally-offset from the excitatory center during outward motion compared to inward motion (Figure S5a).

**Fig. 5.**
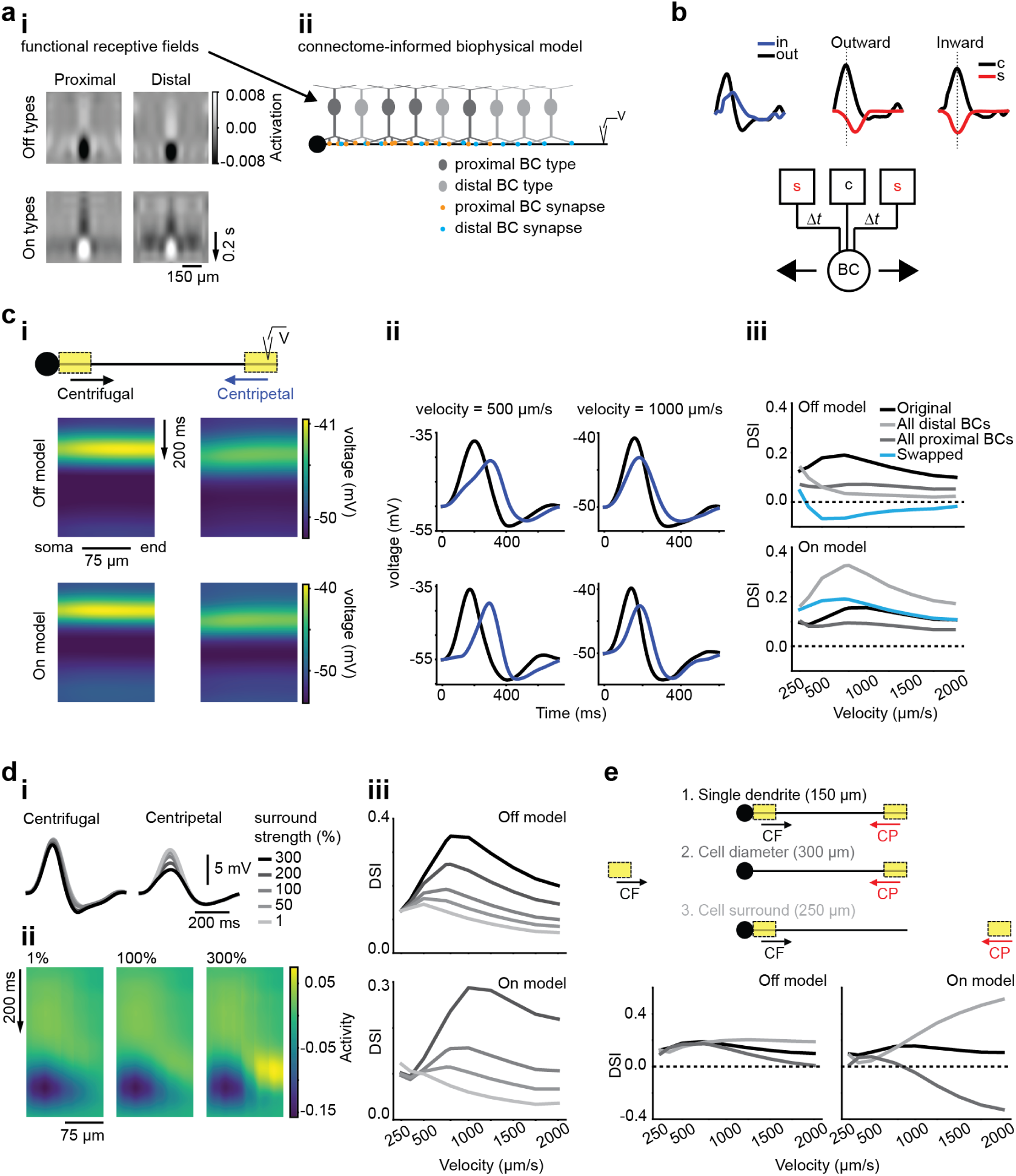
Model amacrine cell inherits bipolar cell motion sensitivity. **(a)** Construction of a model SAC dendrite with accurate BC input. (i) Up-sampled functional RFs of four BC clusters selected based on co-stratification with SAC-connected BC types (**Fig. 4B, Figure S5a, Figure S5b**). The model took inputs from either two Off BC clusters (top) or two On BC clusters (bottom). (ii) Ball-and-stick multi-compartment model of a single SAC dendrite (150 μm long, diagram depicts On model) with BC inputs organized according to anatomical and physiological data (19, 31, 33, 57). **(b)** Modeled response of one up-sampled BC cluster (Off type Proximal) to outward and inward motion stimulation as in (**Fig. 3**). Responses were decomposed into BC RF center (black) and surround (red) contributions (see Figure S5a and Methods for details). These coincide during inward motion, leading to a smaller activation according to a Barlow-Levick detector (circuit, bottom) (54). c, center; s, surround; BC, bipolar cell output; Δt, temporal offset. **(c)** Responses of SAC model to moving bar stimulation in two directions. (i) Membrane potential along the dendrite in an Off (top) and On (bottom) SAC model during CF and CP motion of a moving bar (20 μm) at 1,000 μm/s. (ii) Simulated responses of a distal compartment of the two SAC models for two stimulus velocities (500 and 1,000 μm/s) during the motion. (iii) Directional tuning (DSI) of the distal compartments of the Off and On SAC model dendrites (top and bottom) at different stimulus velocities. We modeled four different BC input distributions. (1) “Original” (black) has BC inputs set based on anatomical and physiological data. (2) “All proximal” (dark gray) replaces distal BC inputs with proximal, for one functional type at all input positions. (3) “All distal” (light gray) replaces proximal BC inputs with distal (4) “Swapped” (blue) assigns functional RFs of the proximal BC type to distal locations and vice versa. (d) Manipulations of surround strength for the SAC model. (i) Off SAC responses to a moving bar (1,000 μm/s) for models using BC RFs with different surround strengths. (ii) Superposition of all BC RFs presynaptic to the Off model at their respective positions along the dendrite at increasing BC surround strengths (from left to right: 1, 100, and 300%). (iii) Directional tuning of the Off and On SAC dendrite models with input from BCs with different RF surround strengths. **(e)** SAC motion dependence on spatial extent of motion stimulus. Top: Different motion stimuli used in the simulations: (1) “Single dendrite”: The bar moves along the 150 μm SAC dendrite. (2) “Cell diameter”: The motion path includes an additional 150 μm extension to the left, where the other side of the cell would be located. (3) “Cell surround”: The motion path contains the SAC dendrite and a 100 μm extension to the right of the dendrite, so that the bar moves off the end of the dendrite. Bottom: Directional tuning of the Off and On SAC model for the different motion stimuli. See also Figure S5a, Figure S5b.

We next asked how the BC clusters extracted based on their RFs correspond to known anatomical BC types. We compared the distribution of IPL depths for ROIs within each cluster to the distribution of BCs in types identified from electron microscopy (EM, data from (18, 19, 31)) and found that the number and extent of co-stratifying clusters was correlated with the number and extent of anatomical BC types (**Fig. 4B**). For instance, we observed three clusters co-stratifying with the stratification band for the three BC types 3a, 3b, and 4. In addition, some of our BC clusters showed a strong correlation with single EM clusters (type 6, type 7). Thus, we argue that these clusters represent distinct types of BCs.

Next, we explored how RF features and motion sensitivity map onto IPL stratification and anatomical type (**Fig. 4C-E**). Within groups of co-stratifying BC types, we found a diversity of RF properties and motion sensing capabilities. Notably, we observed that at least one type within each sublamina of the IPL exhibited motion sensitivity (**Fig. 4E**), suggesting that this functional response property is accessible to post-synaptic partners throughout the IPL. Together, these results suggest that one important aspect of BC diversity is the specification of motion and non-motion signals, a phenomenon observed throughout the mammalian visual system (i.e. in mouse (64–66)).

### Model amacrine cell inherits bipolar cell motion sensitivity

Given the presence of motion sensing BCs throughout the IPL, we wondered whether BCs’ post-synaptic partners use this information for motion sensing. To explore this issue, we constructed biophysical models of On and Off SAC dendrites, which were shown to display a preference for motion from their soma to the distal dendrites (25, 67). Our models were based on previous SAC models and used published connectomic and physiological data about the BC types and locations of BC inputs (33, 57) (**Fig. 5**). But where previous models included none of the center-surround dynamics of the BC RFs, we modeled these spatio-temporal dynamics using RFs derived from specific BC clusters (**Fig. 3**), selecting cluster RFs that were likely matches with BC types known to provide input to SACs (**Fig. 5A**) (see Figure S5a and Figure S5b for details). We observed that many of the BC types chosen for model input were direction selective in our simulations, and further noted that their DS derived from center-surround interactions resembling a Barlow-Levick detector (**Fig. 5A**).

To examine the DS of our SAC models, we simulated a moving bar stimulus that traversed the length of the dendrite in the centrifugal (from soma to distal dendrite, CF) or centripetal (from distal dendrite to soma, CP) direction. We monitored the voltage along the entire model dendrite and found that the model responded with asymmetric depolarization with a preference for CF motion (**Fig. 5B**). In distal model compartments, where SACs have their output synapses, the difference between CF and CP motion was particularly pronounced, and we observed DS across a wide range of physiologically and behaviorally-relevant velocities (3, 4, 64, 66, 68, 69). Previous research on connectomic reconstructions of the SAC has suggested that the gradient of BC types along the SAC dendrite plays a role in their DS (19, 31, 32). We explored this issue by changing which functional RF clusters provide input to our models. We found that some BC cluster RFs led to stronger DS in the SAC model, while others produced weaker DS (**Fig. 5C** and Figure S5b). Thus, BC RFs appear to contribute to SAC DS, and this contribution depends on BC input identity and RF properties.

Next, we explored the influence of the BC RF properties on the DS observed in our SAC models by testing variations in BC surround strength and different spatial stimulation. We tested both weaker and stronger surrounds, particularly because our method of obtaining RFs likely underestimates surround strength (see (70) and **Methods**) and because surround strength can be dynamically altered by environmental conditions (71). First, we changed the strength of the surround component of the BC cluster RFs (Figure S5a). We observed that the model responses to motion were strongly influenced by surround strength, especially in the CP motion direction, presenting a marked surround strength dependence of DS tuning across stimulus velocities (**Fig. 5D**). In addition, we found that the RF surround was the dominant feature conferring DS to the SAC models (Figure S5b). We then evaluated how this surround dependence affects SAC model responses to stimuli traversing different spatial locations and distances relative to the dendrite. In particular, we tested if stimuli that stimulate the proximal BC RF inputs more symmetrically, and thereby reduce the individual BC motion sensitivity (**Fig. 2**), produce less DS in the SAC (“Cell diameter”). Also, we tested if stimuli that activate more of the surround of the distal inputs (“Cell surround”) produces stronger DS. We found that spatial stimulation indeed produced these effects in the model SAC dendrites, especially at high velocities (**Fig. 5E**). Thus, the BC type-specific surround properties play an important role in establishing directional tuning and spatial RF properties of SACs.

### Bipolar cell inputs onto starburst amacrine cells are motion-sensitive

Our modeling results suggest that SACs receive motion-sensitive input from BCs. We confirmed this finding experimentally by imaging glutamate release onto On layer SAC dendrites, by targeted expression of flex-iGluSnFR under control of the ChAT promoter, in response to moving bar (**Fig. 6**) and noise stimuli (**Fig. 7**). With moving bars, we observed a preference for motion originating in the RF center and moving out of the FOV (**Fig. 6F**, *d*′ = 19.4 ± 22.5), a similar pattern of motion sensitivity as we described for iGluSnFR expressed throughout the IPL (**Figs. 1, 2**). In addition, we observed that a moving bar originating outside the FOV and moving through and out again elicited symmetric responses to the two motion directions (**Fig. 6E**; 150 μm, *d*′ = 31.1 ± 20.1; 300 μm. *d*′ = −1.1 ± 18.2). Last, we observed a preference for looming motion compared to receding motion in SAC-layer glutamate release (Figure S3a). These results confirm that motion-sensitive BC clusters provide glutamatergic input to SACs.

**Fig. 6.**
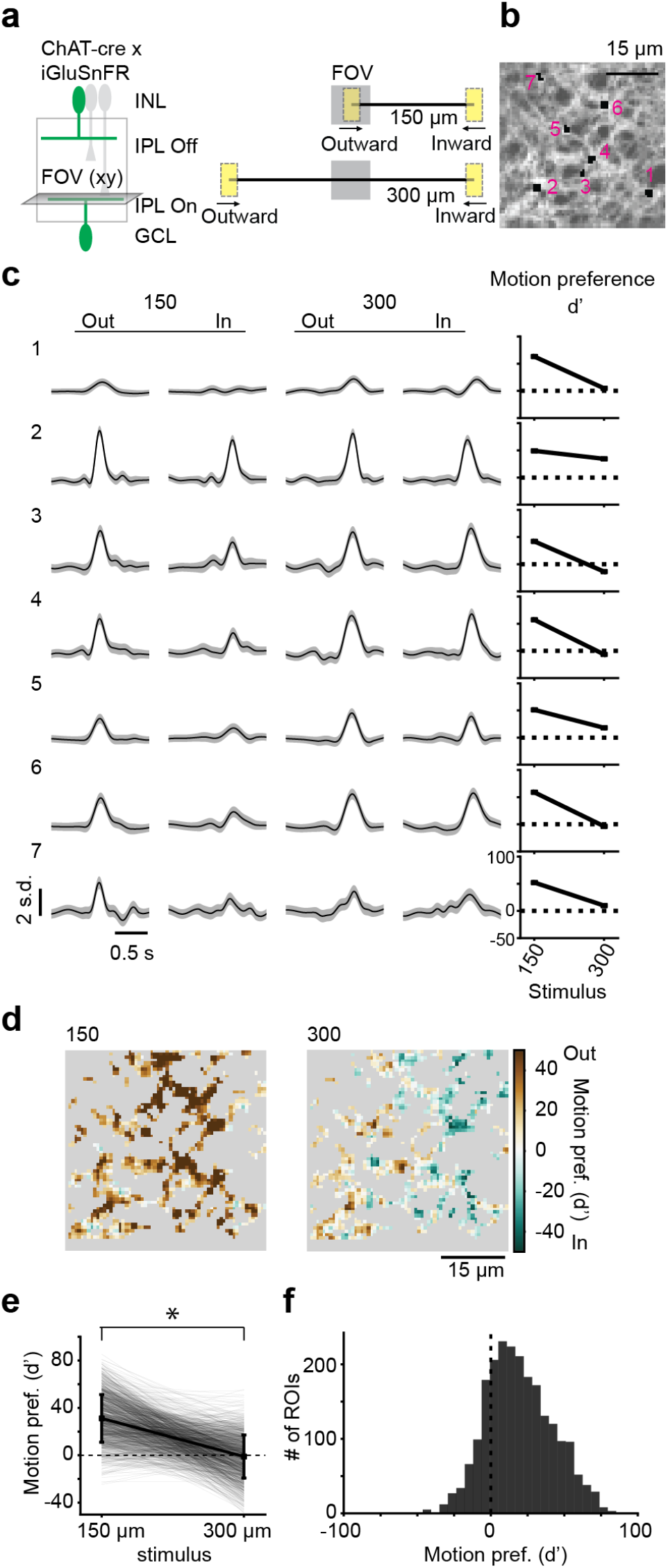
Bipolar cell inputs onto starburst amacrine cells are motion-sensitive. **(a)** Left: flex-iGluSnFR injected into ChAT-cre mice to achieve SAC-specific labeling. Right: ‘Small bar’ stimulus (20 × 40 μm rectangle) moving at 500 μm/s over a distance of 150 or 300 μm, beginning either in the center of the FOV or outside the FOV. Diagram to scale. **(b)** S.d. of the scan field showing iGluSnFR expression in SACs. Black regions/numbers: ROIs in (c). **(c)** Responses predicted with Gaussian Process for each stimulus condition from (a). Numbers correspond to ROIs in (b). Grey shading is 3 s.d. Rightmost column: motion preference (*d*′) for each stimulus travel distance. **(d)** The motion preference (*d*′) for all ROIs in example field for each stimulus travel distance. **(e)** Comparison of motion preference for each stimulus travel distance for all ROIs in the example field (n = 1,134 ROIs). Significant with *p* < 0.001, Wilcoxon test, two-sided. **(f)** Distribution of motion preference for 2,225 ROIs from 2 mice for the 150 μm stimulus distance. Significant with *p* < 0.001, one sample t-test.

**Fig. 7.**
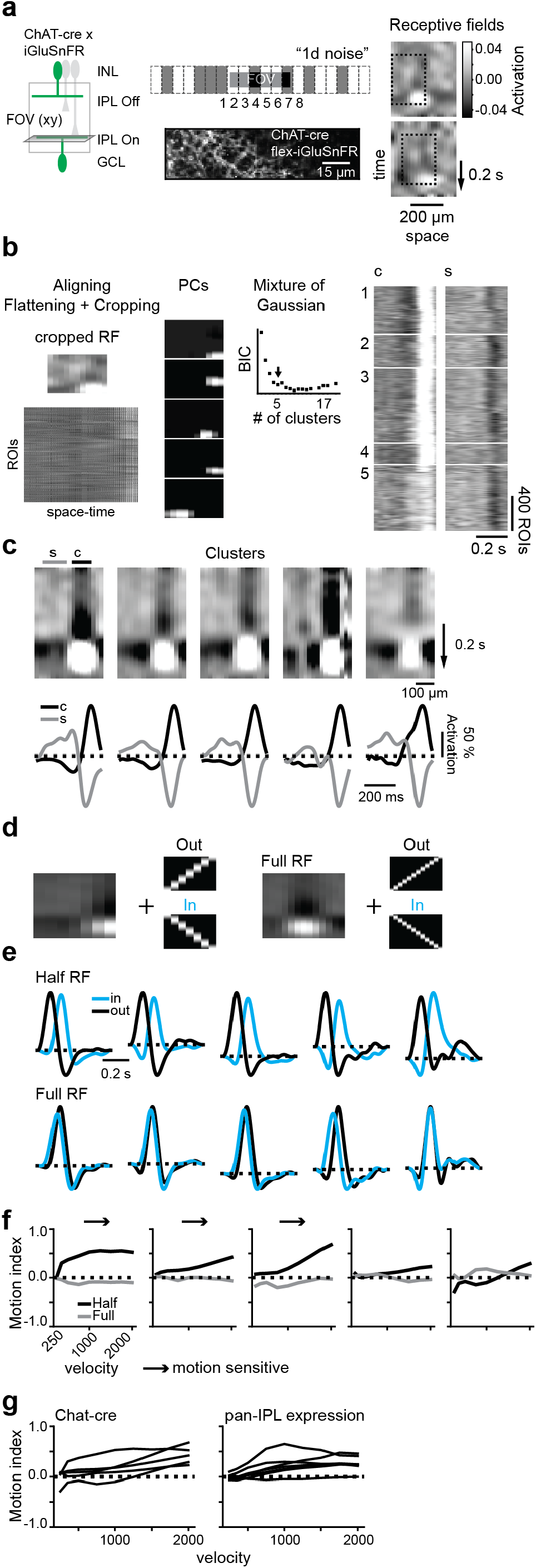
Receptive fields of bipolar inputs onto starburst amacrine cells have diverse motion sensitivity. **(a)** Left: flex-iGluSnFR injected into ChAT-cre mice to achieve SAC specific labeling. Middle: “1D noise stimulus” (top) and s.d. image of FOV (bottom) used to measure RFs from On layer SACs. Right: RFs for two example ROIs. **(b)** Procedure for performing clustering of RFs similar to Fig. 3. Here, the IPL depth is not included as a feature and the optimal number of clusters was 5. Right: center and surround responses of individual ROIs in each cluster. This data set includes 2,725 ROIs from 2 mice. **(c)** Top: Average RFs for each of 5 clusters. Bottom: Average temporal RFs taken from the center (“c”, black) and surround (“s”, gray) regions, normalized to their peaks. **(d)** Convolution with half of the RF or the full RF to measure motion sensitivity, **(e)** Modeled responses to motion (velocity 1,000 μm/s) in two directions for each cluster stimulating the full receptive field or just half of it. **(f)** Motion index as a function of velocity for each cluster for the full (gray) vs half (black) conditions. **(g)** Comparison of velocity tuning curves for On BC clusters identified from mice expressing iGluSnFR only in the SACs (left, ChAT-cre) vs. ubiquitously (right, from Fig. 3).

As further confirmation, we measured and clustered the RFs of BC glutamate release onto On layer SACs (**Fig. 7**). Un-like our findings in **Figure 3**, we selected only five clusters of RFs, which is close to the number of BC types observed to synapse onto On-layer SACs from EM data (4 BC types, (19, 57)) (**Fig. 7B**). We tested the motion sensitivity of these clusters and found that all but one of them exhibited motion sensitivity (**Fig. 7D-F**) and that the clusters’ velocity tuning covered a similar range of motion sensitivity as the On BC clusters uncovered from measuring glutamate release across the IPL (**Fig. 7G**). All together, these results indicate that BC glutamate release onto SACs is motion-sensitive, and that in some stimulus conditions, this asymmetric glutamate release could contribute to motion computations in this amacrine cell type.

### Starburst amacrine cells respond strongly to motion restricted to short distances

Our SAC dendrite model demonstrates a preference for motion restricted to short distances due to the center-surround interactions at the level of the BC inputs (**Fig. 5**). We sought to confirm this stimulus preference through RF mapping of the SAC dendrites. We performed 2-photon Ca^2+^ imaging in a mouse expressing the fluorescent Ca^2+^ sensor flex-GCaMP6f under the control of the ChAT promoter and presented a noise stimulus to map the RF along one axis (**Fig. 8A**). We uncovered RFs of small, varicosity-sized ROIs that exhibited a marked motion preference, and clustered these RFs into groups using Mixture of Gaussian clustering (**Fig. 8B**). These clusters contained ROIs from areas throughout the FOV, and some clusters appeared to contain ROIs from single dendrites (**Fig. 8C**). The average RFs from these clusters revealed three patterns: preferring leftward motion, preferring rightward motion and no motion preference (**Fig. 8D**). These patterns were expected based on the known distribution and outward motion preference of SAC dendrites in the retina (25). We measured the motion trajectory of each ROI’s RF and used this to estimate the preferred motion distance (delta distance) and velocity (**Fig. 8E-F**). We found that many ROIs did not exhibit a motion preference (delta distance near zero) most likely because these ROIs’ dendrites were off-axis from our stimulus. Among motion-preferring ROIs, the preferred motion travel distance peaked at about 70-100 μm, similar to the size of the SAC excitatory RF radius (33) and the estimated motion distance preference from our model (**Fig. 5**). We confirmed that SACs respond in a direction selective manner to a stimulus traveling 100 μm by measuring Ca^2+^ responses in their dendrites to moving bars, and found reliably direction selective responses to this stimulus (**Fig. 8G-K**) with a preference for motion into the FOV (in), as we would expect given the stimulus (**Fig. 8G**). Thus, SAC RFs and direction selectivity appears to be shaped by the motion sensitivity of BCs.

**Fig. 8.**
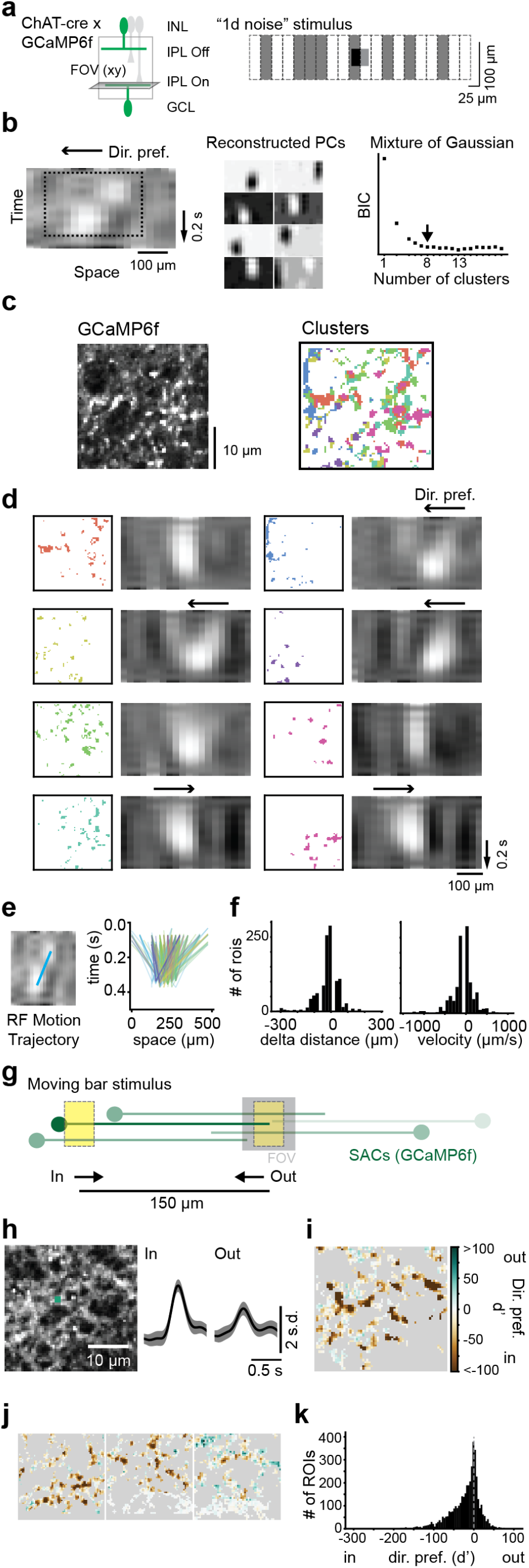
Starburst amacrine cells respond strongly to motion restricted to short distances. **(a)** Left: flex-GCaMP6f mice were crossed with ChAT-cre mice to achieve SAC specific labeling. Right: “1D noise” stimulus presented to SACs expressing GCaMP6f. **(b)** Procedure for clustering ROIs into groups based on their RFs. Left: Example RF that shows a leftward direction preference. Dotted line is the cropped region used for clustering. Middle: Reconstructed components of the sparse PCA. Right: Plot of BIC for different number of clusters using Mixture of Gaussian clustering. Arrow: chosen number of clusters (8). **(c)** Left: s.d. image of a scan field showing GCaMP6f expression in SACs. Right: ROIs color-coded by clusters determined in (b). **(d)** ROIs within each cluster with their average RFs. **(e)** Motion trajectory of the RFs for individual ROIs. Left: example ROI RF showing the estimated motion trajectory (blue line). Right: motion trajectories for all ROIs color-coded by cluster (n=1,112 ROIs, 1 mouse). **(f)** Left: Histogram of the change in center position over time (delta distance) for ROIs from (e). Right: histogram of the RF velocity measured from line slopes in (e). **(g)** Moving bar stimulus (20 × 30 μm bar moving at 500 μm/s) traveling in two directions, either into (“in”) or out of (“out”) the FOV and traversing a distance of 150 μm. Positions of different SACs diagrammed over the stimulus (green dendrites and somas), demonstrating that SACs on the left of the FOV will be optimally stimulated compared to SACs on the right. **(h)** Left: Example scan field (s.d. image) showing GCaMP6f expression and example ROI (green). Right: the Gaussian Process prediction for the response to each motion direction for the example ROI. **(i)** Direction preference (*d*′) estimated from Gaussian Process predictions for all ROIs in the example field in (h). **(j)** Additional fields in this data set showing each ROIs’ direction preference. **(k)** Histogram of direction preference for all ROIs (4,096 ROIs/ 4 fields/ 3 eyes/ 2 mice). Significant with *p* < 0.001, one sample t-test.

## Discussion

In this study, we addressed the question of how BCs in the mouse retina respond to local motion stimulation. We found that some mouse BC axon terminals are sensitive to local motion (**Fig. 1**), responding more strongly to motion originating in their RF center compared to motion originating in their RF surround and traveling to the center (**Fig. 2**). Notably, some BCs exhibit motion sensitivity, while others do not (**Fig. 3**), and the level of motion sensitivity depends on the BCs’ surround strength and timing (**Fig. 4A**). At every depth of the IPL, at least some terminals exhibited motion-sensitive RFs (**Fig. 4B-E**), suggesting that motion signals are available to many different circuits. To determine how BC motion signals are integrated by their postsynaptic partners, we modeled SACs and found that this cell type inherits BC motion signals (**Fig. 5**). We then confirmed that SACs receive motion-selective BC input and showed that their RFs are shaped by the motion sensing properties of BCs (**Fig. 6,7,8**). Our findings suggest that motion signaling arises earlier in the retina than previously thought and that motion vs. non-motion is an important functional distinction between BC types that informs their contribution to retinal processing.

### Bipolar cell motion sensing

In this study, we found that some types of BCs are capable of signaling information about the direction of locally moving objects as well as whether objects are looming or receding. Whether or not BCs transmit this information is highly sensitive to the location of the stimulus relative to the BC’s RF, as well as the cell’s RF properties. Most studies that have previously examined the responses of BCs to moving stimuli did not observe any direction selective tuning in BC membrane potential (72), intra-cellular Ca^2+^ (44), or glutamate responses (42, 43) (but see (46)). All of these studies used global motion stimuli, like gratings or wide moving bars, that originated in the RF surround or outside of the RF of the recorded BCs. Under those stimulus conditions, our modeling predicts that BCs respond symmetrically to stimulation (**Fig. 3**), just as those studies observed. There is one recent study, however, that does not fit into this pattern: using glutamate imaging, they provided evidence for a specialized circuit that bestows “true” DS on a subset of axon terminals in type 2 and 7 BCs (46). Critically, that study found a contribution from wide-field ACs; thus, the mechanism is likely only engaged for large moving stimuli. We found that surround properties were equivalent on two sides of the BC RFs (**Fig. 4**), suggesting that the BC center-surround motion detector operates symmetrically. Thus, the motion detectors described here are not direction selective *per se* but can become so by virtue of their perspective on a motion stimulus.

We found that BC terminals have diverse RF properties that may map onto distinct BC types, including striking differences in the strength and temporal properties of the RF surround that contribute to different motion sensitivity (**Fig. 4**). Many studies have described differences between the RFs of distinct BC types, including differences in the extent and strength of the BC surround (16, 44, 51). RF features are tuned by multiple mechanisms in both the outer and inner retina, with differences in dendritic and axonal spread (18), cone inputs (73), horizontal cell influence (74–76), connectivity to ACs (18), inhibitory receptor complement (53, 77, 78), and susceptibility to neuromodulators (71) all playing a role. How these different factors contribute to the motion sensitivity of BC RFs remains to be determined, though a study published in parallel to ours found that ACs seem not to be necessary to establish motion sensing (79). One key feature aligned with motion sensitivity is a surround that is temporally offset (and slower) than the RF center (**Figs. 4, 5**). Thus, it is possible that an initial center-surround structure established in the outer retina is further fine-tuned in the inner retina, utilizing BC type specific slow AC contributions (i.e. through feedback) to establish strong motion sensitivity.

### Bipolar cell receptive field properties

We found that differences in spatio-temporal RF properties are capable of supporting BC feature selectivity with regard to motion stimuli. Importantly, we found this to be the case while modeling the BC responses using linear models based on their measured RFs, which already revealed complex and diverse motion processing across BC clusters (**Fig. 3**). Increasing evidence suggests that BCs respond in a nonlinear manner to some types of stimuli (37, 55, 56, 80–84), and essentially linearly to other types (85). Our direct observations of motion responses in BCs qualitatively match our response predictions from linear modeling (**Figs. 2, 3**). This could be due to the fact that we model responses based on the full spatio-temporal RFs. In some cases, it is possible that observed nonlinearities in retinal neurons may be the result of either assuming that space and time RFs are separable or taking only space or time components of RFs into account in predictions. It will be interesting to further investigate motion processing in a nonlinear context, which might be particularly important for understanding neuronal responses to natural stimuli. In-deed, the study published in parallel to ours has found that the center-surround RFs of BCs supports novel object detection in natural contexts (79).

### Integration of bipolar cell motion information in down-stream circuits

Although few studies have observed BC motion sensitivity, evidence of this feature of BCs is pervasive in the literature in the form of voltage-clamp recordings of glutamatergic synaptic input to RGCs and ACs. In the mouse retina, the glutamatergic input to both SACs and VG-luT3 ACs recorded in response to looming vs. receding stimuli exhibited a strong looming preference (6, 32, 47), and apparent motion stimuli elicited asymmetric glutamatergic inputs in RGCs (55). In the primate retina, glutamatergic inputs to several types of RGCs were demonstrated to exhibit motion sensitivity (81, 86). And in the rabbit retina, local apparent motion elicited asymmetric glutamatergic inputs to direction selective RGCs (40). In some cases, these results may have been related to voltage clamp errors (87), while in others, they have been attributed to gap junctional interactions between BCs (55, 81). Nonetheless, we propose here that, in some cases at least, these results reflect the motion-sensitive responses we found in subsets of BCs, and that BC RFs play an important role in generating DS, looming sensitivity, and other types of local motion sensitivity through the collection of BC inputs in diverse downstream neurons.

We explored how BC motion sensitivity contributes to one downstream motion computation, DS in SACs. In our measurements of BC glutamate release onto SAC dendrites, we observed clear DS during local motion stimulation starting in BC RF centers. Our modeling suggests that these direction selective inputs are integrated to support SAC DS, and are in line with SACs’ selectivity for motion towards their dendritic tips (25). Many studies that have previously evaluated SAC DS used stimuli that activate BC RFs’ center-surround motion detector, including local moving bars (33), differential motion stimuli (88), and expanding rings (25,67, 89,90). We argue that this stimulus dependence is not a bug but a feature of SAC RFs, tuning them to prefer local motion starting close to the SAC soma. Stimulus-dependent motion processing has previously been described in mouse VGluT3-expressing ACs (91), W3 RGCs (68), and rabbit On-Off direction selective RGCs (92), all of which show preferences for local motion. In addition, directional tuning in some On-Off direction selective RGCs in mouse is stronger for local drifting gratings compared to global ones (88) and in rabbit directional tuning in direction selective RGCs is observed for stimuli traveling distances shorter than the spacing between photoreceptors (93). On the other hand, direction selective RGCs are known to play important roles in brain functions and behaviors involving global motion information (64, 94, 95). Previous studies have also suggested that BCs participate in direction detection via other mechanisms (for example see (19, 31, 32, 35, 46, 57) but also (33, 34, 42)) which raises the question of how these mechanisms work together. Given the diverse stimuli often used to probe motion processing, it is possible that distinct mechanisms of direction detection are engaged under different environmental conditions (as previously suggested in (96, 97)), which could ensure robust DS. Thus, studying the role of BC motion signals during local motion processing in SACs and direction selective RGCs could provide important insights to understand the role of BCs in this circuit. Notably, the mechanism of signal suppression during null direction motion that we report here has long been described (54) and has also been observed in the fly visual system (reviewed in (98)) and in the rodent whisker system (99).

Beyond the DS circuit, there are many other RGC types that could rely on BC motion sensitivity information. In mammals, several RGC and AC types are object motion sensitive, responding specifically to local motion or differential motion (4, 6, 15, 68, 72, 85, 91, 100–102), including several prominent primate RGC types, such as parasol RGCs (81, 86). In general, RGCs and ACs must fulfill a few requirements to be capable of integrating BC motion information into their computations. The first requirement is that downstream cells receive input from motion-sensitive BC types. It will be interesting to explore the wiring of BCs to their postsynaptic partners in this context. The second requirement is that down-stream cells should employ post-synaptic integration that allows for motion-sensitive information to be preserved in the cell’s output. RGCs are capable of retaining RF structure from BCs (103); indeed, center-surround interactions at the level of BCs contribute to RGC encoding of spatial features (56, 82). A mechanism for local motion integration is hinted at by a recent study that found that the dendrites of some mouse RGC types perform less spatial averaging than others (104). Since spatial averaging would likely blur spatially-restricted local motion signals (**Fig. 2**), this integration feature could allow for integration of motion information from BCs. Combining connectomic information about wiring with functional and modeling explorations of RGC and AC responses that take BC RF properties into account, such as we have done here, will thus provide a fruitful avenue for understanding motion processing in the retina.

The BC motion sensitivity observed here may be relevant in a wide variety of natural conditions important for behavior. Local object motion and looming detection are highly relevant to animals (reviewed in (105, 106)) and are highly salient to humans (107–109). In the case of the mouse, they represent prey and predators, respectively (3–6). The BC motion detector is particularly primed to detect moving objects that are initially occluded in a scene, such as a grasshopper jumping out of the grass or a hawk diving from a great distance. At the same time, the BC motion detector is rather insensitive to the type of scene motion that occurs when the body, head and eyes smoothly move. This dichotomy allows for detection of behaviorally-relevant moving objects (15). Thus, it is striking that this essential visual information for animal survival is detected already in bipolar cells.

## ACKNOWLEDGEMENTS

We thank Gordon Eske, Merle,, Ilona Los, and Zhijian Zhao for excellent technical support, Robert G. Smith and Gregory Schwartz for feedback and discussions, and all members of the Euler, Berens, and Franke labs for technical and conceptual feedback. Funding was provided by the German Research Foundation (DFG BE 5601/2-1; SPP 2041 with BE 5601/4-1 and EU 42/9-1; EXC 2064 ML, project number 390727645), the German Ministry of Education and Research (Bernstein Award (01GQ1601) to PB, the Bernstein Center for Computational Neuroscience (01GQ1002) to KF, the Tübingen AI Center to PB (01IS18039A), Christiane Nuesslein-Volhard-Stiftung to AV, Universitätsklinikum Tübingen Fortüne Fellowship to MK and AV, the European Research Council (ERC-StG “NeuroVisEco” 677687) to TB, The Wellcome Trust (Investigator Award in Science 220277/Z20/Z) to TB, the UKRI (BBSRC, BB/R014817/1) to TB, and the Max Planck Society (M.FE.A.KYBE0004) to KF.

## Author contributions

(CRediT taxonomy, https://jats4r.org/credit-taxonomy): Conceptualization: SS, TB, PB, TE, AV; Data curation: MK, AV; Formal analysis: SS, AV; Funding acquisition: TB, PB, TE, AV; Investigation: MK, AV, TS, KF; Methodology: SS, AV, KF, YR, TB, TE, PB; Project administration: PB, TE, TS; Resources: TE, PB; Software: TE, AV, SS; Supervision: TE, PB, AV, KF, YR; Validation: AV, SS; Visualization: AV, SS; Writing - original draft: AV, SS, MK; Writing - review and editing: all authors

## Materials and Methods

### Animal and tissue preparation

All animal procedures were approved by the governmental review board (Regierungspräsidium Tübingen, Baden-Württemberg, Konrad-Adenauer-Str. 20, 72072 Tübingen, Germany) and performed according to the laws governing animal experimentation issued by the German Government. For measuring glutamate release in the IPL, we used either the ChAT-cre transgenic line (n = 3; JAX 006410, The Jackson Laboratory; (110)) or C57Bl/6 J (n = 4, JAX 000664) mice. For Ca^2+^ imaging in SACs, the ChAT-cre transgenic line was crossbred with the Cre-dependent green fluorescent reporter line Ai59D (n = 3; JAX 024105; (111)). We used adult mice greater than 6 weeks old of either sex. Owing to the exploratory nature of our study, we did not use randomization and blinding. No statistical methods were used to predetermine sample size. Animals were housed under a standard 12 h day/night rhythm at 22°and 55% humidity. For activity recordings, animals were dark-adapted for >1h, then anesthetized with isoflurane (Baxter) and euthanized by cervical dislocation. The eyes were enucleated and hemisected in carboxygenated (95% O_2_, 5% CO_2_) artificial cerebrospinal fluid (ACSF) solution containing (in mM): 125 NaCl, 2.5 KCl, 2 CaCl_2_, 1 MgCl_2_, 1.25 NaH_2_PO_4_, 26 NaHCO_3_, 20 glucose, and 0.5 L-glutamine (pH 7.4). Throughout the experiments, the tissue was continuously perfused with carboxygenated ACSF at ~36°C, containing ~0.1 μM Sulforhodamine-101 (SR101, Invitrogen) to reveal blood vessels and any damaged cells in the red fluorescence channel (112). All procedures were carried out under very dim red (>650 nm) light. The positions of the fields relative to the optic nerve were not taken into account in this study. In some cases the retina was cut into pieces and each piece was mounted and imaged separately to prolong the light sensitivity of the tissue.

### Virus injection

We injected the viral constructs AAV2.7m8.hSyn.iGluSnFR (generated in the Dalkara lab - for details, see (113); the plasmid construct was provided by J. Marvin and L. Looger (Janelia Research Campus, USA)) or AAV9.CAG.Flex.iGluSnFR.WPRE.SV40 (Penn Vector Core) into C57Bl/6 J and ChAT-cre mouse lines, respectively. A volume of 1 μL of the viral construct was injected into the vitreous humour of 4 to 6-week-old mice anesthetized with 10% ketamine (Bela-Pharm GmbH & Co. KG) and 2% xylazine (Rompun, Bayer Vital GmbH) in 0.9% NaCl (Fresenius). For the injections, we used a micromanipulator (World Precision Instruments) and a Hamilton injection system (syringe: 7634-01, needles: 207434, point style 3, length 51 mm, Hamilton Messtechnik GmbH). Imaging experiments were performed 3-4 weeks after injection.

### Two-photon imaging

We used a MOM-type two-photon microscope (designed by W. Denk, MPI, Heidelberg; purchased from Sutter Instruments/Science Products; (112)). As described before, the system was equipped with a mode-locked Ti:Sapphire laser tuned to 927 nm (MaiTai-HP DeepSee, Newport Spectra-Physics), two fluorescence detection channels for iGluSnFR/GCaMP6f (HQ 510/84, AHF/Chroma) and SR101 (HQ 610/75, AHF), and a water immersion objective (W Plan-Apochromat ×20 /1.0 DIC M27, Zeiss). For image acquisition, we used custom-made software (ScanM by M. Müller and T.E.) running under IGOR Pro 6.37 for Windows (Wavemetrics), taking time-lapsed 64 × 64 pixel image scans (at 9.766 Hz) or 128 × 32 pixel image scans (at 15.625 Hz). For vertical glutamate imaging in the IPL, we recorded time-lapsed 64 × 56 pixel image scans (at 11.16 Hz) using an electrically tunable lens (ETL; for details, see (62)).

### Light stimulation

A DLP projector (lightcrafter (LCr), DPM-E4500UVBGMKII, EKB Technologies Ltd) with internal UV and green light-emitting diodes (LEDs) was focused through the objective. The LEDs were band-pass filtered (390/576 Dualband, F59-003, AHF/Chroma), for spectral separation of the mouse M- and S-opsins, and synchronized with the microscope’s scan retrace.

In our experiments, photoisomerization rates ranged from ~0.5 (black image) to ~20 × 10^3^ P* per s per cone for M- and S-opsins, respectively (for details, see (114)). Two-photon excitation of photopigments caused additional steady illumination of ~10^4^ P* per s per cone (discussed in (112, 115, 116)). The center of the light stimulus was adjusted to be on the center of the recording field, and was verified post-hoc either using the receptive fields (RFs) measured from noise or by estimating the location at which the stimulus response onset was fastest for moving bar stimuli. Analysis was adjusted if the stimulus was determined to be off center. For all experiments, the tissue was kept at a constant mean stimulator intensity level for at least 15 s after the laser scanning started and before light stimuli were presented. Stimuli were presented using custom Python software (QDSpy: https://github.com/eulerlab/QDSpy).

Four types of light stimuli were used: (*i*) small, positive contrast moving bar (20 × 40 μm for iGluSnFR, 20 × 30 μm for GCaMP6f) appearing in different locations relative to the FOV and moving at 500 μm/s over varied distances (appearance locations, motion directions, and distances specified in **Figures 1, 2, 6, 8** with 2-3 s between each stimulus presentation; (*ii*)a “long bar with aperture” stimulus with moving bar (40 × 385 μm) traveling in two directions at 500 μm/s through a small aperture box (80 μm^2^) (**Fig. 1**). (*iii*) “1-d noise stimulus” consisting of 20 adjacent rectangles (20 × 50 μm for iGluSnFR, 25 × 100 μm for GCaMP6f), with each rectangle independently presenting a random black and white (100% contrast) sequence at 20 Hz for 2.5-5.0 s (**Figs 3, 4, 7**). (iv) a white looming and receding stimuli consisting of a white spot on black background that appeared and then expanded or retracted at a velocity of 800 μm/s. For looming, the spot started at 10 μm and expanded to 600 μm (Figure S3a). All stimuli were presented at 100% contrast, were presented in the same pseudo-random order for each imaging field, and were achromatic, with matched photoisomerization rates for mouse M- and S-opsins.

### Data analysis

Data analysis was performed using Python 3 and IGOR Pro. Data were organized in a custom-written database schema using DataJoint for Python framework (https://datajoint.io/) (117).

### Pre-processing

Pre-processing was performed using custom scripts in IGOR Pro and Python. First, we measured the s.d. of each pixel and discarded the bottom 50-90% from further analysis. The threshold depended on the experiment: for ubiquitously expressing iGluSnFR, 50-70% of pixels were discarded; for ChAT-cre restricted imaging, 70-90% were discarded because fewer pixels in the imaging field exhibited fluorescence. Traces for each remaining pixel were imported into DataJoint, then high-pass filtered using a Butter filter (0.2 Hz, order = 5) and z-normalized by subtracting each traces’ mean and dividing by its s.d. A stimulus time marker embedded in the recorded data served to align each pixel’s trace to the visual stimulus with 1.6 - 2 ms precision. For this, the timing for each pixel relative to the stimulus was corrected for sub-frame time-offsets related to the scanning.

### Motion sensitivity estimation

To measure average responses of ROIs during low resolution iGluSnFR imaging (**Fig. 1**, Figure S3a), we drew manual rectangular ROIs at different locations relative to the stimulus position and calculated a binned average of the ROIs’ pixels’ responses, resampling the response times of each pixel to 63 Hz. This allowed us to resolve higher time resolution than the frame frequency of our imaging and retain the precise alignment to the stimulus timing.

To obtain Gaussian Process (GP) estimates for BC glutamate release and SAC dendritic Ca^2+^, we followed the methods in (59). First, pixel response quality was assessed by calculating the response quality index (as in (16)) for each stimulus condition separately. Pixels were discarded if the stimulus condition with the largest quality index value fell below 0.35. Then, ROIs were built automatically from each high quality pixel to include neighboring high quality pixels and to have dimensions around 2 μm (3-9 pixels, average ROI diameter in **Fig. 2**: 2.29 ± 0.28*μm;* in **Fig. 6**: 2.03 ± 0.34*μm;* in **Fig. 8I-K**: 1.41 ±0.20*μm*), which is the estimated size of BC boutons (16) and near the resolution limit of our imaging. ROIs were allowed to have some overlap with one another, which improved the signal to noise of our models and made no assumptions about the resolution of our imaging. Because of this, maps of d-prime (*d*)′ in **Figures 2D, 6D, 8I-J** report the measured value only at the center pixel of a ROI. The average response of a ROI’s pixels was obtained by resampling the response times at 125 Hz and averaging within time bins.

Then, for each ROI, we created a GP estimate of the response trace using the GPy toolbox (https://sheffieldml.github.io/GPy) at 50 Hz, with warping of the time resolution during the period when the moving bar was presented to capture fast response kinetics (59). For a given ROI, all stimulus conditions were included in the model. We used the Sparse Gaussian Process Regression algorithm with the Radial Basis Function kernel (with parameters with kernel variance = 1.1 and kernel lengthscale = 0.05), and then the model prediction was stored in DataJoint. GP estimates whose mean activity had an s.d. below 0.1 across time for all stimulus conditions were discarded from further analysis, as these regions were considered non-responsive.

*d*′ was estimated for each ROI’s GP estimate as follows: the peak response (*μ*) and the s.d. at this peak (*σ*) during the time of bar presentation was measured. For each pair of opposite directions, *d*‣ was calculated as:

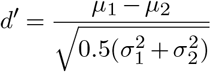

For each imaging field, the location of the FOV relative to the stimulus was assessed based on RF mapping (see below) if available or based on the relative response timing of stimuli in opposite directions.

### Receptive field mapping and clustering

RFs were obtained using a modified spike-triggered averaging method that employs a spline basis to estimate smooth RFs (RFEst toolbox: https://github.com/berenslab/RFEst, (63)). First, traces for each pixel and the stimulus trace were up-sampled to the scan line precision (1.6-2 ms) using linear interpolation to align stimuli and responses. The stimulus trace was then mean subtracted so that 50% contrast was set to zero. Then, we formed ROIs using the same method as described for Gaussian Process ROIs (above) except that we did not discard low quality pixels before creating ROIs. ROI diameters in **Fig. 3**: 2.04 ± 0.07*μm;* in **Fig. 7**: 2.69 ± 0.33*μm;* in **Fig. 8b-f**: 1.38 ± 0.21*μm*. To restrict ROIs to the IPL in X-Z recordings using the ETL, the border lines of the IPL were manually determined using an s.d. image. Then, we utilized the splineLG function from RFest to obtain the smoothed spike-triggered average for each ROI over a time lag of 0.5 s.

To cluster RFs, we performed sparse principal components analysis (PCA) and mixture of Gaussian (MoG) clustering using libraries and custom scripts in Python as follows: First, we aligned the RF center for each ROI to the same spatial position. Due to the noisy nature of the individual ROI RFs, we accomplished this by first clustering the ROIs within a field using a hierarchical clustering algorithm (SciPy cluster.hierarchy.linkage in Python, https://www.scipy.org, (118)(119)) and grouping ROIs into clusters using a fixed distance criterion (0.05). This allowed us to obtain average RFs for ROIs with similar RFs within a field, which had the same RF center and polarity. We measured the maximum in these cluster averages (or minimum for Off layer ROIs) and defined this as the RF center for all ROIs in the cluster.

Next, RFs for all ROIs were flattened to one dimension (space-time) and cropped to include the region of the RF that was available for all ROIs. At the precision of our stimulus alignment, it was possible for the stimulus to be off-center of the imaging FOV by up to 100 μm, resulting in a shift of the mapped RFs and in our data set an over-representation of one half of the RF (see **Fig. 3** and Figure S3b). Thus, clustering was performed on just half of the RF. Next sparse principal components (PCs) of the flattened RFs were determined using the sparsePCA function (120) from scikit-learn (https://scikit-learn.org, (121)). We also determined the depth of each ROI’s center in the IPL using the manually-determined IPL boundaries to find the percentage of the IPL thickness at the ROI’s center. Together, the RF sparse PCs and IPL depth constituted the feature weights for MoG clustering, which was performed using the scikit-learn mixture.GaussianMixture toolkit (121). To determine the best number of clusters, we varied the targeted number of clusters between 3 and 19 and estimated the Bayesian information criterion (BIC). Next, we calculated the average RF for each cluster and estimated the temporal kernels for center and surround in distinct spatial regions from these averages. We measured several parameters from these temporal kernels: (*i*) latency was the time between the peak of the center and peak of the surround response; (*ii*) surround strength was measured as the ratio of the surround peak and center peak; (*iii*) biphasic index was measured by finding the ratio of the maximum and minimum of the center’s temporal kernel; (*iv*) center full-width half maximum (FWHM) was determined by calculating the mean spatial kernel during the time of the center response, fitting this to a Gaussian and finding the FWHM of that Gaussian. The clustering procedure was performed separately for each of the data sets in **Figures 3, 7, 8**. Anatomical correlation between the clusters found in **Fig. 3** and BC types identified from previously published electron microscopy (EM) reconstructions (18) was performed by obtaining the kernel density estimation using Gaussian kernels (KDE, scipy.stats.gaussian_kde) of the IPL depth of the ROIs in each cluster. These KDE curves were correlated with each BC type from EM to determine the stratification overlap (**Fig. 4**).

### Statistical testing

Statistical testing was performed using Python packages pingouin (122) for Wilcoxon test (**Figs. 2,6**) and SciPy’s stats package (118) for 1 sample t-test (**Fig. 8**), Spearman correlation coefficient (**Fig. 4**), and paired t-test (Figure S2b).

### Modeling bipolar cell responses from receptive fields

To predict BC responses to moving stimuli from their RFs, we performed convolution between the RFs and stimulus images. The RFs were cropped to contain the full RF or just half of the RF to model responses to different stimulation in space. Convolution was performed at each spatial location of the images independently, and then summed across space to obtain the final temporal predictions of the responses. We determined the direction selective or looming selective index by measuring the peak response during stimulation in each direction:

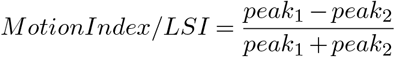

To prepare cluster RFs for use in the SAC biophysical model, we increased the RF resolution in both space and time, denoised the RFs, and used only half of the RF, reflected, to create BC model inputs Figure S5a. To maintain the space-time structure of the RF during interpolation, we performed singular value decomposition (SVD), performed linear interpolation to increase the resolution of the space and time components by 20x and 1.6x, respectively, and then reconstructed the space-time RFs from the first three components. This denoised the RFs while increasing their resolution. Finally, we mirrored the RF to create a full (symmetrical) spatial RF.

To manipulate the strength of the surround in **Fig. 5**, Figure S3a, and Figure S5a we selected values of opposite polarity to the RF center (negative values for On, positive values for Off) in the surround. These values were multiplied by a scalar (0.01, 0.5, 2, 3) to increase or decrease the strength of the surround. For Off RFs, we found that the surround was generally weaker. This could be due to two features of our “1D noise” stimulus - first, that the background on which the row of rectangles was presented was dark, suppressing the surround of Off BCs, and second, that the individual rectangles of the stimulus were small, leading to low total contrast of the stimulus, which has been demonstrated to cause underestimates of surround strength (70). Thus, we tested the larger increase in surround strength of 300% specifically for Off RFs. In contrast, with 300% surround, On BC cluster surrounds were so strong that resulting model responses were completely suppressed (data not shown).

### Starburst amacrine cell model

To design the anatomical distribution of BC input to the SAC model dendrite, we calculated the number of BC synapses in 10 μm dendritic segments from glutamatergic input labeling in SAC dendrites (33) and assigned them to BC types according to anatomical data about BC type-specific wiring to SAC dendrites (57). The Off model included anatomical types 1 and 3a, the On model included anatomical type 7 and a generic type 5 (by merging types 5o, 5t and 5i into one). The BCs’ RFs were represented by the functional RFs at their respective locations (**Figs. 3, 4**). Where multiple BC clusters overlapped, we tested each possible RF cluster (Figure S5b). Moving bar stimuli and BC responses were calculated as described above. We included a spontaneous baseline BC activity, which could be regulated up or down by stimulation of the BC center and surround, respectively. BC activity was rectified by clipping values below zero. The BC activation across time became the current injection input to the SAC model dendrite at the respective synapse locations of each BC. The input to each model was scaled such that the maximum depolarization in the most distal model compartment would reach approximately −35 mV at the lowest stimulus velocity. The biophysical SAC ball-and-stick model was implemented in Brian2 (https://brian2.readthedocs.io, (123)). The multicompartment model consisted of an iso-potential soma (diameter: 7 μm) and a 150 μm long dendrite. The initial 10 μm of the dendrite had a diameter of 0.4 μm, the remaining dendrite had a diameter of 0.2 μm (33). In addition to a leak current, the model included Ca^2+^ channels in the distal third of the dendrite (124). The calcium current was translated to a change in the calcium concentration via γ_*Ca*_^2+^ and the Ca^2+^ concentration in each compartment decayed according to an exponential model (125, 126) with time constant τ_*Ca*_^2+^ (See **Table 1**). The strength of tuning in the SAC was measured in the distal third of the dendrite. We calculated the DSI from the membrane potential for each compartment in the distal dendrite from the response to centrifugal (CF) and centripetal (CP) motion as

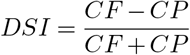

and reported the average of those compartments in the velocity tuning curves (**Fig. 5** and (Figure S5b)).

**Table 1.** SAC model parameters.

| Table 1. SAC model parameters |  |  |  |
| --- | --- | --- | --- |
| Parameter | Value (unit) | Parameter | Value (unit) |
| $R_i$ | 150 $\Omega \cdot \text{cm}$ (33) | $E_{\text{Ca}^{2+}}$ | 120.0 mV (127) |
| $R_m$ | 21,700 $\Omega \cdot \text{cm}^2$ (33) | $\bar{g}_{\text{Ca}^{2+}}$ | 0.013 mS/mm<sup>2</sup> (127) |
| $C_m$ | 1 $\mu\text{F}/\text{cm}^2$ (33) | $[\text{Ca}^{2+}]_0$ | 50 nM (125) |
| $E_L$ | -54.4 mV (127) | $\tau_{\text{Ca}^{2+}}$ | 5 ms (125) |
| dt Brian2 | 0.1 ms | $\gamma_{\text{Ca}^{2+}}$ | 20 M/nC |

### Code and data availability

Data as well as Python code will be available upon publication from our GitHub repository https://github.com/eulerlab/bc-motion and http://www.retinal-functomics.org.

