## Supplemental Figures for "Center-surround interactions underlie bipolar cell motion sensing in the mouse retina"

Figure S2a: related to Figure 2
Figure S2b: related to Figure 2
Figure S3a: related to Figure 3
Figure S3b: related to Figure 3
Figure S5a: related to Figure 5
Figure S5b: related to Figure 5

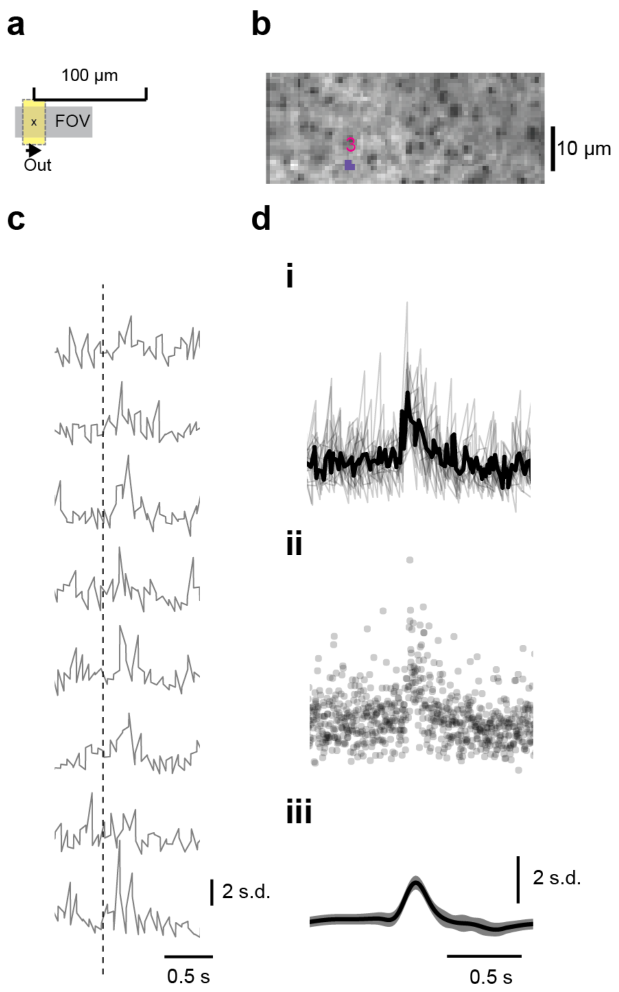


**Figure S2a. Gaussian Process modeling of BC terminal responses, related to Figure 2.**
(a) The moving bar stimulus as in **Fig. 2**.
(b) ROI 3 from **Fig. 2**
(c) Z-scored example single trials in response to the stimulus in (a).
(d) Comparison of averaging and Gaussian Process prediction. (i) Single trials (gray) overlaid with the binned average taken at 63 Hz. (ii) Scatter plot of all measurements from all trials. (iii) Gaussian process prediction. Grey shading is 3 s.d. around the estimated mean.

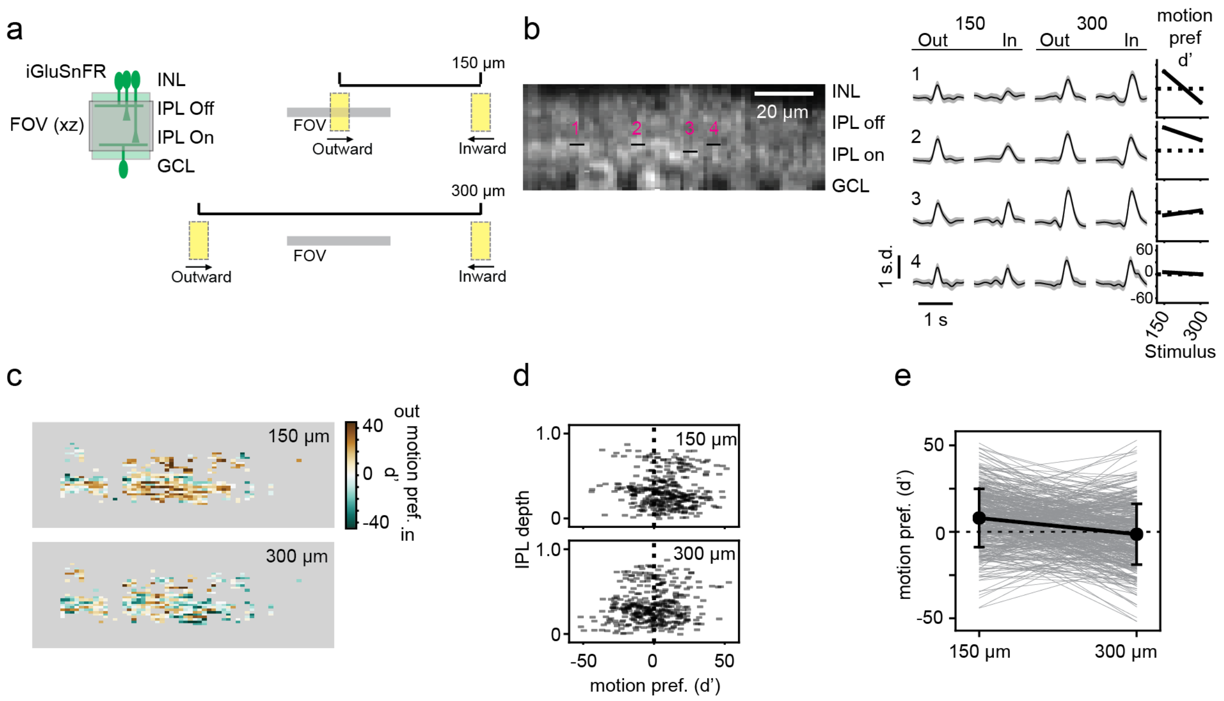


**Figure S2b. Bipolar cell responses to moving bar stimuli across the inner plexiform layer, related to Figure 2.**

(a) Left: iGluSnFR is ubiquitously expressed in retinal neurons, including in the cells of the inner plexiform layer (IPL, green region). INL, inner nuclear layer; GCL, ganglion cell layer; FOV, field of view. Right: Moving bars (20 x 40 µm) presented to the retina traveling in two directions (out vs. in) and traversing two distances (150, 300 µm). All objects to scale.

(b) Example ROIs (black regions, numbered) in the On layer of the IPL overlaid with s.d. of the imaged field and their responses to the stimuli in (a) as predicted using Gaussian Process modeling. Grey shading is 3 s.d. Rightmost column: motion preference ($d'$) for each stimulus travel distance. Positive values represent a preference for motion in the "out" direction.

(c) The motion preference ($d'$) for all ROIs in the field.

(d) ($d'$) as a function of IPL depth for all ROIs in the field.

(e) Motion preference ($d'$) for each ROI in the population for each stimulus condition. The two conditions are significantly different (150 µm, $d' = 8.3 \pm 16.9$; 300 µm, $d' = -1.5 \pm 17.5$; paired T-test, $p < 0.01$). Sample size in is 381 ROIs/ 1 field/ 1 mouse.

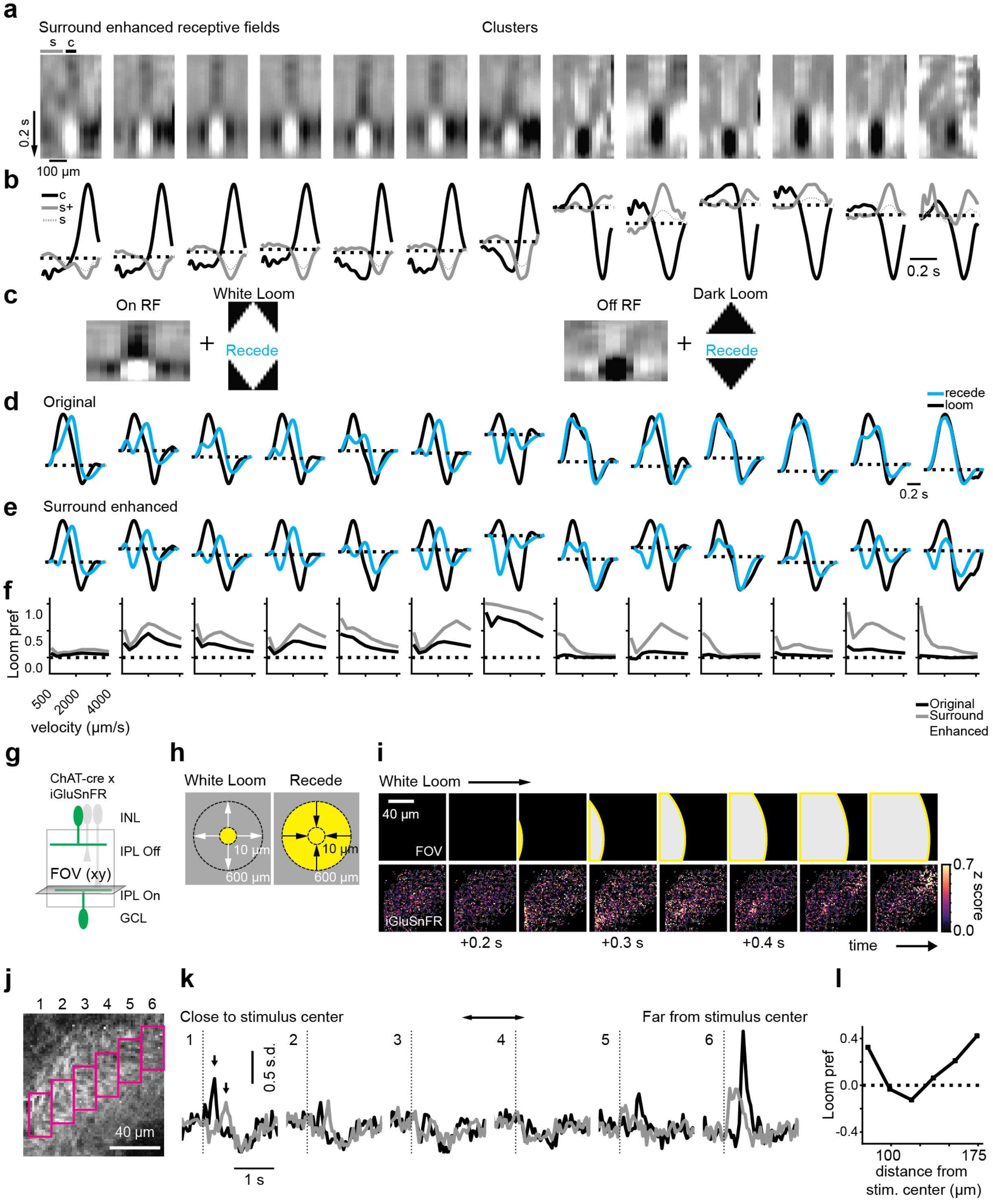


**Figure S3a. Modeling reveals differing looming preference of bipolar cell clusters, related to Figure 3.**
(a) Cluster average RFs with the surround strength enhanced. On BCs were enhanced 150%, while Off BCs were enhanced 300%. “c” and “s” show the regions used to calculate the spatial average of the center and surround in (b).
(b) Average temporal RFs taken from the center (“c”) and surround (“s”) regions indicated in (a), showing the relative amplitude of center and surround. For surround, dotted (“s”) = original surround, solid (“s+”) = enhanced surround.
(c) Example RF convolved with looming and receding stimuli to model responses of BC clusters.
(d) Example modeled responses of each BC cluster to looming and receding stimuli at a stimulus velocity of 1,000 µm/s using the cluster average RFs from **Figure 3**.
(e) Example modeled responses of each BC cluster to looming and receding stimuli at a stimulus velocity of 1,000 µm/s using

the surround enhanced RFs in (a).
(f) Looming preference of each BC cluster's RF across stimulus velocities. Looming-sensitive BC clusters exhibit even stronger looming preference when the surround is strengthened.
(g) Experiment to validate looming response predictions. flex-iGluSnFR was injected into ChAT-cre mice to achieve SAC-specific labeling.
(h) White looming and receding stimuli. For looming, a white spot appears on a dark background with a diameter of 10 µm and expands to 600 µm at a rate of expansion of 800 µm/s, then disappears. Receding stimulus is the reverse in time. Diagram not to scale.
(i) Response of one FOV to looming stimulus. Top row: position of the stimulus in the FOV. The center of the stimulus is outside the FOV. Bottom row: montage of the average z-scored fluorescence response of glutamate sensor iGluSnFR during stimulation.
(j) Average iGluSnFR fluorescence during stimulation, showing six ROIs used to measure fluorescence responses.
(k) Mean binned fluorescence in response to each stimulus condition for the pixels in each ROI. Arrow show peak responses used to calculate the looming sensitivity index (LSI).
(l) LSI for the responses in (k) vs. the ROI distance from the stimulus center. The position closest to the center exhibits looming sensitivity predicted by our modeling in (d)-(e).

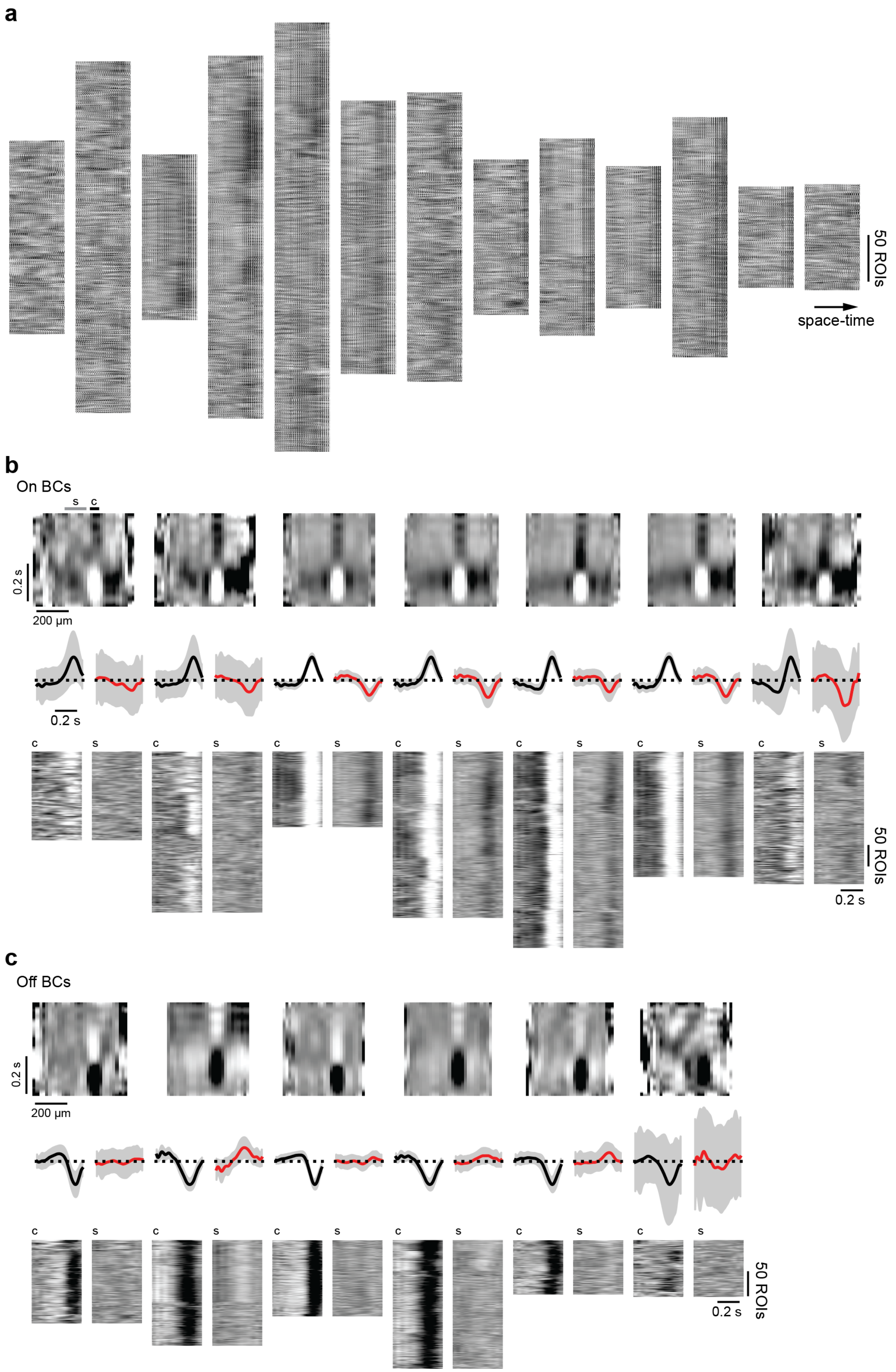


**Figure S3b. Clustered bipolar cell receptive fields and in-cluster variability, related to Figure 3**
(a) Images of individual ROIs' RFs, cropped and flattened in space-time, sorted by cluster. Clusters are ordered by IPL depth.
(b) RFs of On BC clusters. First row: average receptive field of each cluster, showing additional edge regions that were not

used for modeling. Data is noisier because sample sizes are lower for more peripheral regions. “c” is center region used for extracting time kernels, “s” is surround region. Second row: center (black) and surround (red) time kernels for each cluster, normalized to the peak of the center response. Grey is s.d. Third row: center (“c”) and surround (“s”) time kernels for each individual ROI, sorted by cluster.
(c) RFs of Off BC clusters, conventions as in (b).

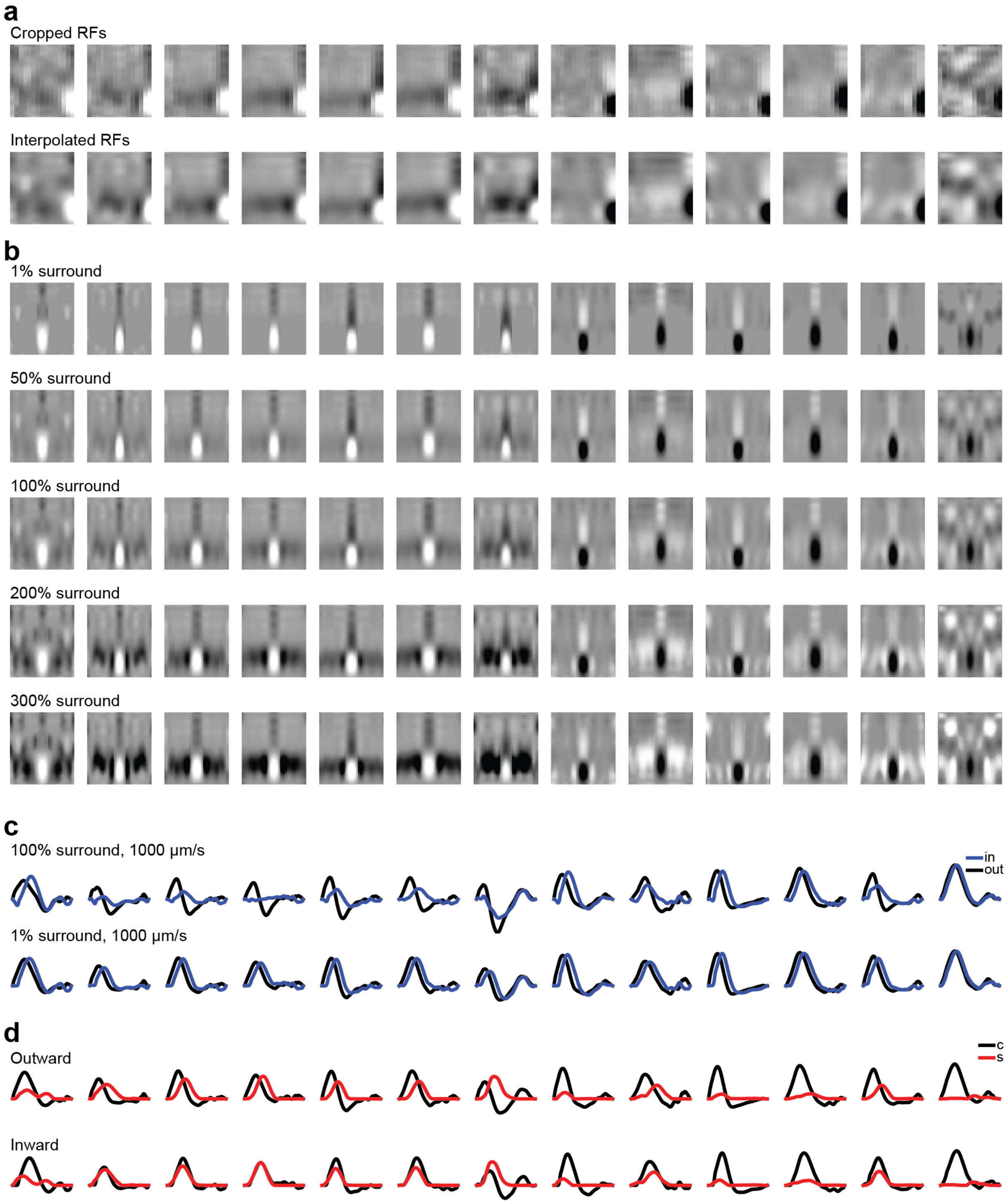


**Figure S5a. Receptive field modification for biophysical modeling and center vs. surround timing across clusters, related to Figure 5.**

(a) Top: Original average RFs from clusters in **Fig. 3** were cropped to include the more-complete half of the RF. Order of clusters is the same as in **Fig. 3** - ordered based on average IPL depth from GCL to INL. Bottom: For each RF, we performed singular value decomposition (SVD). Then, we performed linear interpolation on spatial and temporal components of the SVD. Finally, the RFs were reconstructed from the first 3 spatial and temporal components to create the RFs shown.

(b) RFs prepared for biophysical SAC modeling are shown with different levels of surround strength. The RFs in (a) were reflected to create a full RF for each cluster. To change the strength of the surround, we multiplied all values that were opposite polarity to the center (i.e. negative values for On clusters, positive values for Off clusters) in the surround spatial region by a scalar multiplier. This changed the strength of the surround while maintaining the same strength of the center region.

(c) Convolution of 1,000 μm/s moving bar stimulus across half the RF (as in **Fig. 3i**) for the 100% and 1% surround cases from (b) to model responses to moving stimuli. Motion sensitivity is largely lost when the surround is minimized.
(d) Decomposition of modeled responses to outward and inward motion. Center ("c", black) response was estimated as the response to 1% surround shown in (c). The surround ("s", red) contribution was estimated by subtracting the response modeled from RFs with 1% surround from the response predicted from 100% surround RFs. In motion-sensitive clusters, the timing of excitatory center and inhibitory surround are more offset for outward motion than for inward motion due to the center and surrounds' different temporal properties (see also **Fig. 4**).

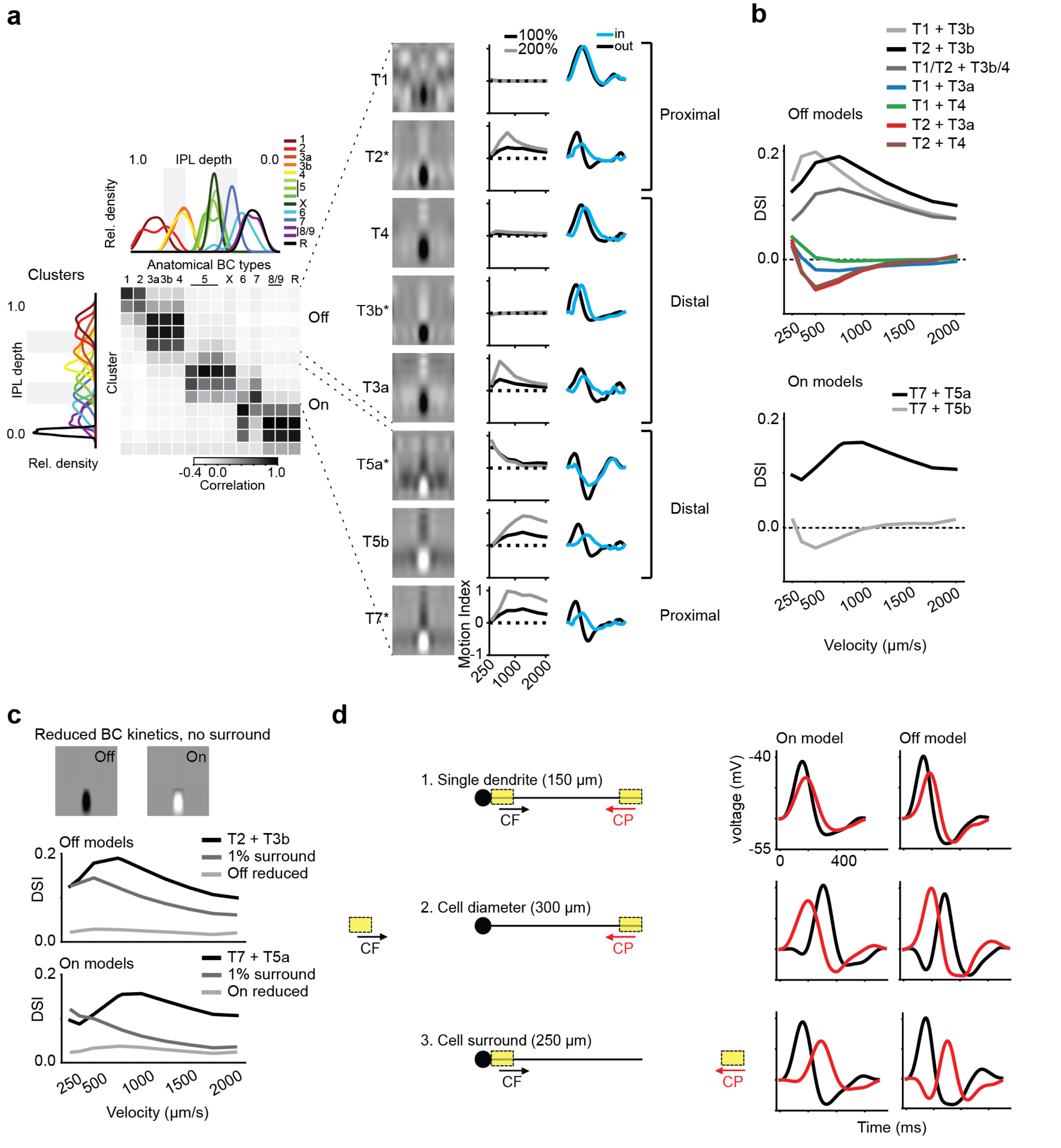


**Figure S5b. SAC models with different functional BC clusters and motion paths, related to Figure 5.**

(a) Left: Pairwise correlation between IPL stratification of BC clusters identified in the current study and anatomically (18, 19, 31) (see also **Fig. 4**). Right: RFs of BC clusters that costratify with SACs (8 out of 13 identified BC clusters), their motion sensitivity across velocities, and their responses to moving bar stimuli at 1,000 µm/s. Types were assigned according to IPL stratification correlation. Location-specific wiring (proximal vs. distal) of BC types along SAC dendrites (19, 31, 57) is indicated on the right. Asterisk indicates the types chosen for further modeling in **Fig. 5**.

(b) Directional tuning of SAC models using different combinations of functional BC types. Due to co-stratification, some type labels of BC clusters might need to be permuted (e.g., BC type 1 and 2, see discussion in (16)). We explored all possible combinations of one distal and one proximal type during modeling. In addition, we constructed an Off model containing four BC types, two proximal and two distal BC types (labeled "T1/T2 + T3b/4"). Our results suggest that placing BC type 3b and

5a in the Off and On models' distal positions, respectively, is crucial for establishing a preference for CF motion. In the Off model, the inclusion of BC type 2 compared to type 1 results in a slight shift of the tuning curve towards higher velocities in the two-type model. The inclusion of further types results in a slight reduction of the DSI at low stimulus velocities. We found that BC type 2, but not type 1, conferred surround dependence of tuning, suggesting that the tuning in the type 1 model relies on a different mechanism (details not shown). Perhaps for type 1 BCs, the interaction of BC types with different temporal receptive field center properties could play a role in the preference for CF motion (19, 31, 57). Taken together, these results indicate that the properties of BC RF types determine the directional tuning of postsynaptic model SAC dendrites.
(c) Comparison of models with and without BC RF components. Top: "Reduced" BC type 2 and 7 RFs. The biphasic center response was manipulated to only include the initial response and all surround activity was removed. Middle and bottom: Comparison of the original Off and On models from **Fig. 5c** (black) with the "1% surround" models from **Fig. 5c**(dark gray) and the models using "reduced" RFs (light gray). Removing BC kinetics resulted in a reduction of the DSI at all stimulus velocities. In line with previous work (33), the SAC models with limited BC kinetics still exhibit a slight preference for CF motion. This behavior likely results from the skewed BC input distribution on the SAC dendrites, which favors sequential activation of inputs in the CF direction (33).
(d) Example depolarizations of the On and OFF SAC models in response to the different spatial stimuli described in **Fig. 5** (Stimulus velocity: 1,000 µm/s). In comparison to the motion stimulus across a single dendrite, the extension of motion to the cell diameter results in a reduction of the preference for CF motion in the On model and even a switch towards a preference for CP motion in the Off model. This is due to the more symmetric activation of the BC RFs. On the other hand, the extension of the motion towards the cell surround resulted in an increase in the preference for CF motion because the surrounds of more BCs are activated in the CP direction. These findings are most prominent at high stimulus velocities (see **Fig. 5**). These results suggest that the BC input leads to a SAC RF that favors local motion stimuli originating near the SAC's soma.
